# AD-linked R47H-*TREM2* mutation induces disease-enhancing proinflammatory microglial states in mice and humans

**DOI:** 10.1101/2020.07.24.218719

**Authors:** Faten A. Sayed, Lay Kodama, Joe C. Udeochu, Li Fan, Gillian K. Carling, David Le, Qingyun Li, Lu Zhou, Hansruedi Mathys, Minghui Wang, Xiang Niu, Linas Mazutis, Xueqiao Jiang, Xueting Wang, Man Ying Wong, Fuying Gao, Maria Telpoukhovskaia, Tara E. Tracy, Georgia Frost, Yungui Zhou, Yaqiao Li, Matthew Brendel, Yue Qiu, Zuolin Cheng, Guoqiang Yu, John Hardy, Giovanni Coppola, Shiaoching Gong, Fei Wang, Michael A. DeTure, Bin Zhang, Lei Xie, Dennis W. Dickson, Wenjie Luo, Li Gan

**Affiliations:** Neuroscience Graduate Program, University of California, San Francisco, San Francisco, CA, USA; Gladstone Institute of Neurological Disease, San Francisco, CA, USA; Helen and Robert Appel Alzheimer’s Disease Research Institute, Brain and Mind Research Institute, Weill Cornell Medicine, New York, NY, USA; Medical Scientist Training Program and Neuroscience Graduate Program, University of California, San Francisco, San Francisco, CA, USA; Department of Neurobiology, Stanford University School of Medicine, Stanford, CA, USA; The Picower Institute for Learning and Memory, Department of Brain and Cognitive Sciences, Massachusetts Institute of Technology, Cambridge, MA, USA; Icahn School of Medicine at Mount Sinai, Department of Genetics and Genomic Sciences, NY, USA; Tri-Institutional Computational Biology & Medicine Program, Weill Cornell Medical College, NY, USA; Computational and Systems Biology Program, Memorial Sloan Kettering Cancer Center, New York, NY, USA; Departments of Psychiatry and Neurology, Semel Institute for Neuroscience and Human Behavior, David Geffen School of Medicine, University of California, Los Angeles, Los Angeles, CA, USA; Chemical Biology Program, Weill Graduate School of Medical Sciences of Cornell University, New York, NY, USA; Institute for Computational Biomedicine, Dept. of Physiology and Biophysics, Weill Cornell Medicine, New York, NY, USA; Department of Computer Science, Hunter College, & The Graduate Center, The City University of New York, New York, NY, USA; Bradley Department of Electrical and Computer Engineering, Virginia Polytechnic Institute and State University, Arlington, VA, USA; Department of Neurodegenerative Disease, UCL Queen Square Institute of Neurology, Queen Square, London, UK; Department of Population Health Sciences, Weill Cornell Medical College. New York, NY, USA; Mayo Clinic, Jacksonville, Florida, USA

## Abstract

The hemizygous R47H variant of *TREM2*, a microglia-specific gene in the brain, increases risk for late-onset Alzheimer’s disease (AD). In this study, we identified a subpopulation of microglia with disease-enhancing proinflammatory signatures associated with the R47H mutation in human AD brains and tauopathy mouse brains. Using transcriptomic analysis at the single-nuclei level from AD patients with the R47H or the common variant (CV*)-TREM2* with matched sex, pathology and *APOE* status, we found that the R47H mutation was associated with cell type- and sex-specific transcriptional changes in human AD brains, with microglia exhibiting the most robust alterations. Further characterization revealed that R47H-associated microglial subpopulations had enhanced inflammatory signatures including hyperactivation of Akt, one of the signaling pathways downstream of TREM2. In a newly-generated tauopathy knock-in mouse model expressing one allele of human *TREM2* (*hTREM2)* with either the R47H mutation or CV, R47H induced and exacerbated tau-mediated spatial memory deficits in female mice. Single-cell transcriptomic analysis of microglia from these mice also revealed transcriptomic changes induced by R47H that had significant overlaps with R47H microglia in human AD brains, including robust increases in proinflammatory cytokines, activation of Syk-Akt-signaling, and elevation of a subset of disease-associated microglial signatures. Strikingly, pharmacological Akt inhibition largely reversed the enhanced inflammatory signatures induced by R47H in primary microglia treated with tau fibrils. By unraveling the disease-enhancing properties of the R47H mutation in mouse and human, our findings shed light on an immune-linked AD subtype and provide new directions for modulating microglial immune responses to treat AD.

## MAIN

Alzheimer’s disease (AD) is the most common form of late-onset dementia. In addition to the pathological hallmarks of amyloid plaques and neurofibrillary tangles composed of hyperphosphorylated tau, AD is also characterized by increased microglial activation and an upregulation of cytokines in the brain. These aberrant microglial phenotypes in AD have been largely considered responses to the toxic buildup of plaques and tangles. However, genome-wide association studies have identified many risk alleles for late-onset sporadic AD that are highly expressed in microglia ^1–4^, providing compelling genetic evidence for important roles of microglia in AD pathogenesis. Among these risk genes, *TREM2* is the strongest immune-specific risk factor identified to date, with the heterozygous R47H point mutation significantly increasing the odds ratio of developing late-onset AD ^1, 2^.

TREM2 is a single transmembrane receptor expressed exclusively in cells of the myeloid lineage, especially microglia ^5, 6^. Upon ligand engagement, TREM2, together with its adaptor DAP12/TYROBP, recruits SYK and triggers several signaling cascades such as PI3K-AKT and MAPK pathways ^7, 8^. These TREM2-dependent pathways in turn regulate many microglial functions, including inflammatory cytokine secretion, proliferation, phagocytosis, cell survival through mTOR signaling, and synapse elimination ^9–20^. While the exact ligands in the brain remain elusive and are likely to be context-dependent, TREM2 binds *in vitro* to apoptotic cells ^11, 18^, anionic ligands ^11, 21^, apolipoproteins ^22–24^, and amyloid *β* ^25–27^. Cleavage of TREM2 by metalloproteinases releases soluble TREM2 ^28–30^, which may regulate microglial cell survival and inflammation ^31, 32^.

Accumulating studies suggest that TREM2 is involved in the stepwise activation of microglia in neurodegenerative processes. In the context of neurodegenerative mouse models, Trem2 is required for the conversion of microglia into disease-associated microglia (DAM) or a microglial neurodegenerative phenotype (MGnD) ^33, 34^. This MGnD microglia-state can be activated by apoptotic cells and is partially mediated through Trem2’s interaction with Apoe ^33^. These microglia are characterized by downregulation of homeostatic genes, such as *P2ry12, Tmem119, Sall1,* and *Cx3cr1,* and upregulation of pro-inflammatory signatures such as *Apoe, Axl, Tlr2, Cd74,* and *Itgax.* Currently, it is unclear whether this DAM state is neuroprotective or neurotoxic for disease progression. Deletion of mouse Trem2 (*mTrem2)* prevents microglial conversion to this disease-state and protects against tau-induced atrophy ^35, 36^. *mTrem2* deficiency in amyloid models, however, leads to increased amyloid toxicity, likely due to the role of Trem2 in plaque compaction ^37–40^. Furthermore, human AD-microglia seem to be enriched in some of these DAM genes, such as *APOE, CD74, HLA-DRB1,* and show overlap in molecular pathways related to lipid and lysosomal biology. However, there is likely to be human-specific AD-microglia subpopulations since many gene signatures do not overlap between the mouse and human AD-associated microglia ^41, 42^. These observations suggest the role of TREM2 and DAMs in neurodegenerative diseases is context- and disease state-specific.

Little is known about how the R47H mutation of *TREM2* contributes to AD. Previous studies reported that AD patients carrying the heterozygous R47H variant show higher neuritic plaque densities, reduced microglial coverage of amyloid plaques and more severe plaque-associated neuritic dystrophy, as well as increased accumulation of autophagosomes in microglia ^9, 40, 43^. Human AD patients carrying the R47H mutation also display higher levels of both total tau and phosphorylated tau in CSF compared to non-carriers ^44, 45^. One transcriptomic study showed that at the bulk-tissue level, several immune-related genes are decreased in R47H carriers such as *IRF8, HLA-DRA* and *AIF1*, suggesting either a decrease in the number of microglia or decreased expression of these genes on a per-cell basis ^46^. However, comprehensive transcriptomic studies at either the single-cell level or with a large sample size of patient brain tissues have not been done. In mouse models, homozygous knock-in of R47H human *TREM2* (R47H-*hTREM2*) on a background lacking endogenous mTrem2 leads to deficits in microglial amyloid plaque compaction and increases tau staining and dystrophic neurites bypassing plaques ^40, 47^. Similar to the *mTrem2* deficiency amyloid mouse model, protein expression of amyloid-plaque-induced microglial activation markers, such as *C1qa, Lyz2,* and *Spp1* is decreased ^46^, suggesting that R47H may be neurotoxic due to its inability to activate microglia and compact amyloid plaques. In a recent study, male PS19 tauopathy mice expressing homozygous R47H-*hTREM2* exhibited reduced tau phosphorylation, brain atrophy, and synapse loss compared to mice expressing CV-*hTREM2* ^48^, similar to the phenotype of *mTrem2*-deficient tauopathy mice. However, how the R47H mutation elevates AD risk in humans remains unclear.

In the current study, we uncovered an R47H-enriched microglia subpopulation in human AD brains by performing single-nuclei RNA sequencing analysis of 16 AD patient brain tissues with and without the TREM2-R47H mutation. To investigate the functional changes induced by R47H in AD, we used CRISPR to replace one allele of *mTrem2* in mice with CV- or R47H-*hTREM2*, generating a heterozygous R47H-*hTREM2* mouse model that was then crossed to a tauopathy model. We showed that in response to tau pathology, R47H-associated microglia upregulated a subset of DAM signatures and increased expression of pro-inflammatory cytokines, which was partially mediated through Akt hyperactivation, one of TREM2’s downstream signaling molecules. Together, our study sheds new light on the disease-enhancing mechanisms of the R47H mutation in AD.

## RESULTS

### R47H Mutation Induces Cell Type and Sex-Specific Transcriptional Changes in Human AD Brain Tissues

To dissect the pathogenic mechanisms associated with *TREM2^R47H^* in human AD patients, we performed single-nuclei RNA-seq (snRNA-seq) of mid-frontal cortical tissues from 16 AD brains that harbored a single allele of the TREM2 common-variant (CV) or the R47H mutation (n=8 per group, Supplementary Table 1). The samples were matched in sex and *APOE E3/E4* genotype (Fig. 1a), as well as age, amyloid and tau burdens (Extended Data Fig. 1a-c). Following an established human snRNA-seq protocol ^41^, we sequenced 111,932 nuclei and used 83,051 nuclei for downstream analysis after removal of potential multiplets using DoubletFinder ^49^ and filtering for low-quality nuclei determined by thresholding gene counts, UMI counts, and percent mitochondrial genes per nuclei (Fig. 1a, Extended Data Fig. 1d-g, Supplementary Table 2). Using reference gene sets for cluster annotations ^50, 51^, we identified the major cell types of the brain and observed that cell types were similarly represented in all samples sequenced (Fig. 1b-c, Extended Data Fig. 1h-i).

**Figure 1.**
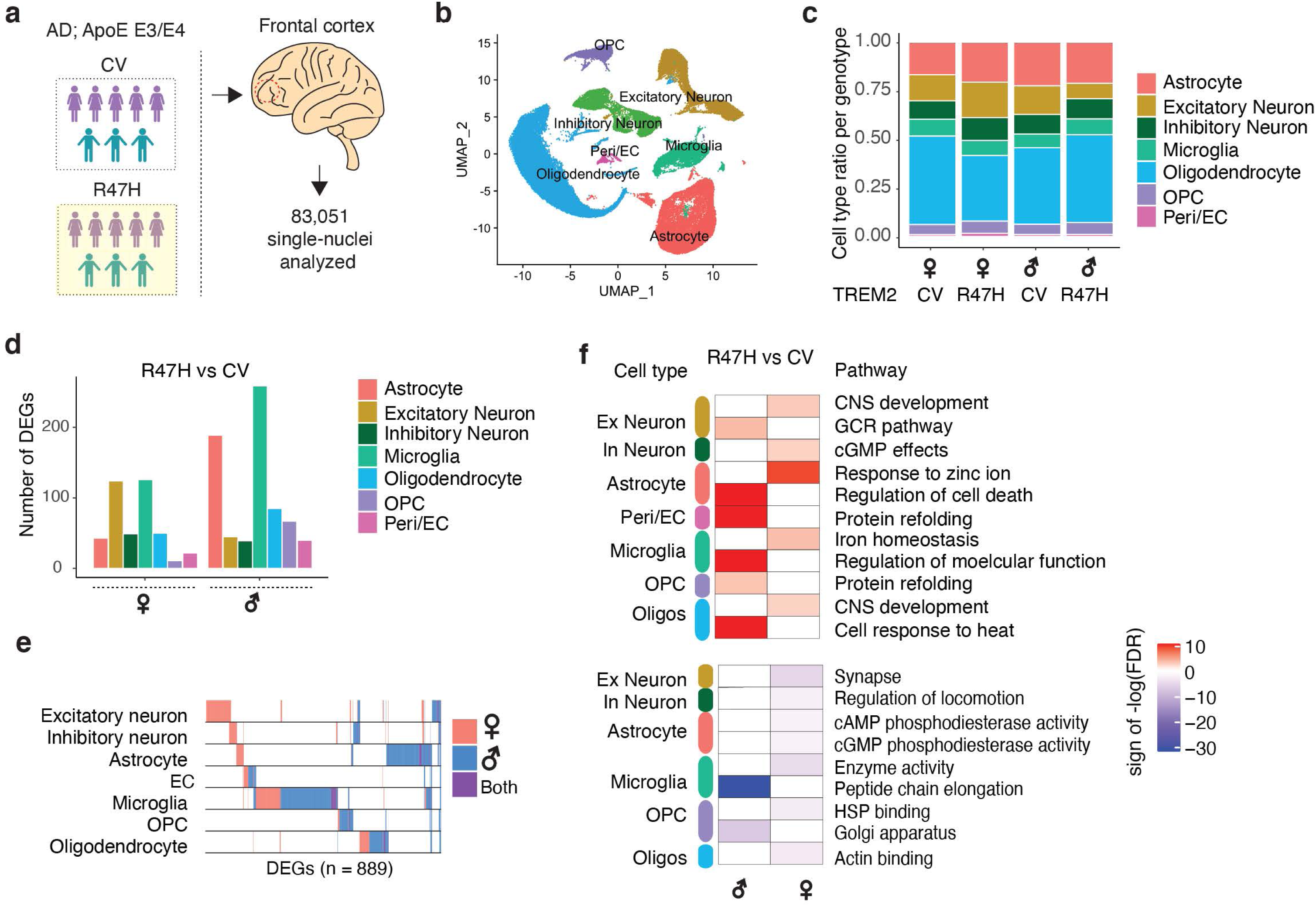
R47H Mutation Induces Cell Type- and Sex-Specific Transcriptional Changes in Human AD Patient Brains. **(a)** Schematic showing the sex and genotypes of age-matched human donors used for single-nuclei RNA-seq. n = 83,051 nuclei were isolated from the mid-frontal cortex of AD patients carrying the CV-TREM2 variant (n=8) and AD patients carrying the R47H-TREM2 variant. Purple and turquoise cartoons denote females and males, respectively. See also Supplementary Table 1. **(b)** UMAP plots of all single nuclei and their annotated cell types. Peri/EC = pericyte/endothelial cells, OPC = Oligodendrocyte precursor cells. **(c)** Proportion of cell types for each of the 4 genotypes. **(d)** Number of DEGs between R47H vs. CV samples for each cell type and each sex. FDR<0.05. **(e)** Binary plot indicating whether a gene (column) is a DEG or not in a given cell type (rows) or in each sex (pink: female; blue: male; purple: overlapping in both sexes) (n=889 DEGs). **(f)** Heatmap of the Gene Ontology pathway enrichment analysis for DEGs determined in (d). Colors denote upregulated (+1, red, upper panel) or downregulated (−1, blue, lower panel) pathway multiplied by the -log10(FDR). Ex Neuron = Excitatory neurons, In Neuron = Inhibitory neurons, Oligos = oligodendrocytes. See also Extended Data Fig. 1 and Supplementary Tables 1-3.

We first performed differential expression analysis to compare the effects of the R47H mutation in each cell type and sex. The mutation was associated with many transcriptional changes in all cell types in both sexes, with microglia having the largest total number of DEGs (Fig. 1d, Supplementary Table 3). Interestingly, *TREM2^R47H^* microglia induced sex-specific transcriptomic changes, with a significantly higher number of DEGs in male compared to female microglia and astrocytes, and far fewer changes in male compared to female excitatory neurons. We then examined the overlaps between the DEG signatures in different cell types and found that few R47H-induced DEGs were shared between male and female brains (purple, Fig. 1e). There was also little overlap of the DEGs among different cell types (rows, Fig. 1e). Indeed, pathway analysis revealed that the top molecular pathways represented by the DEGs were cell-type and sex-specific (Fig. 1f).

### Human R47H AD-Microglia Exhibit Unique Microglial Disease Signatures Characterized by Hyperactivation of Syk-Akt Signaling

To further dissect the transcriptomic changes in microglia induced by *TREM2^R47H^*, we subclustered the 4350 microglia cells from all samples and identified 9 different transcriptional states (Fig. 2a, Supplementary Table 4). We focused on clusters 1-5, as clusters 6-9 had fewer than 30 cells each. When split by *TREM2* genotype, we found differential distributions of microglial subclusters between *TREM2^R47H^* and *TREM2^CV^* samples, with clusters 4 and 5 being more enriched in *TREM2^R47H^* samples (Fig. 2b-c), as well as differentially distributed by sex (Extended Data Fig. 2). Cluster 4 and 5 cells had distinct transcriptomic signatures when compared with homeostatic microglial cluster 1 cells (Fig. 2d,e). We then compared the gene signatures of these five microglial clusters identified in our AD *TREM2^R47H^* transcriptomic dataset with previously-identified microglial signatures associated with high-pathology from AD *TREM2^CV^* brains ^41^ (Mathys-AD, Fig. 2f). Clusters over-represented in the R47H carriers (clusters 4 and 5) had the most overlap, suggesting that *TREM2^R47H^* led to further enrichment of the human AD-associated microglia subpopulations (Extended Data Fig. 3a). One of the upregulated markers in R47H-associated clusters was MS4A6A, an important AD risk gene that regulates soluble TREM2 in human AD ^52^. Other interesting components of the R47H-associated signature were acyl-coA synthase (ACSL1), which is involved in lipid metabolism and inflammation, serglycin (SRGN), an intracellular proteoglycan interacting with inflammatory mediators, and SLC11A1, a divalent transition metal (iron and manganese) transporter with antimicrobial function (Fig. 2d,e, Extended Data Fig. 3a).

**Figure 2.**
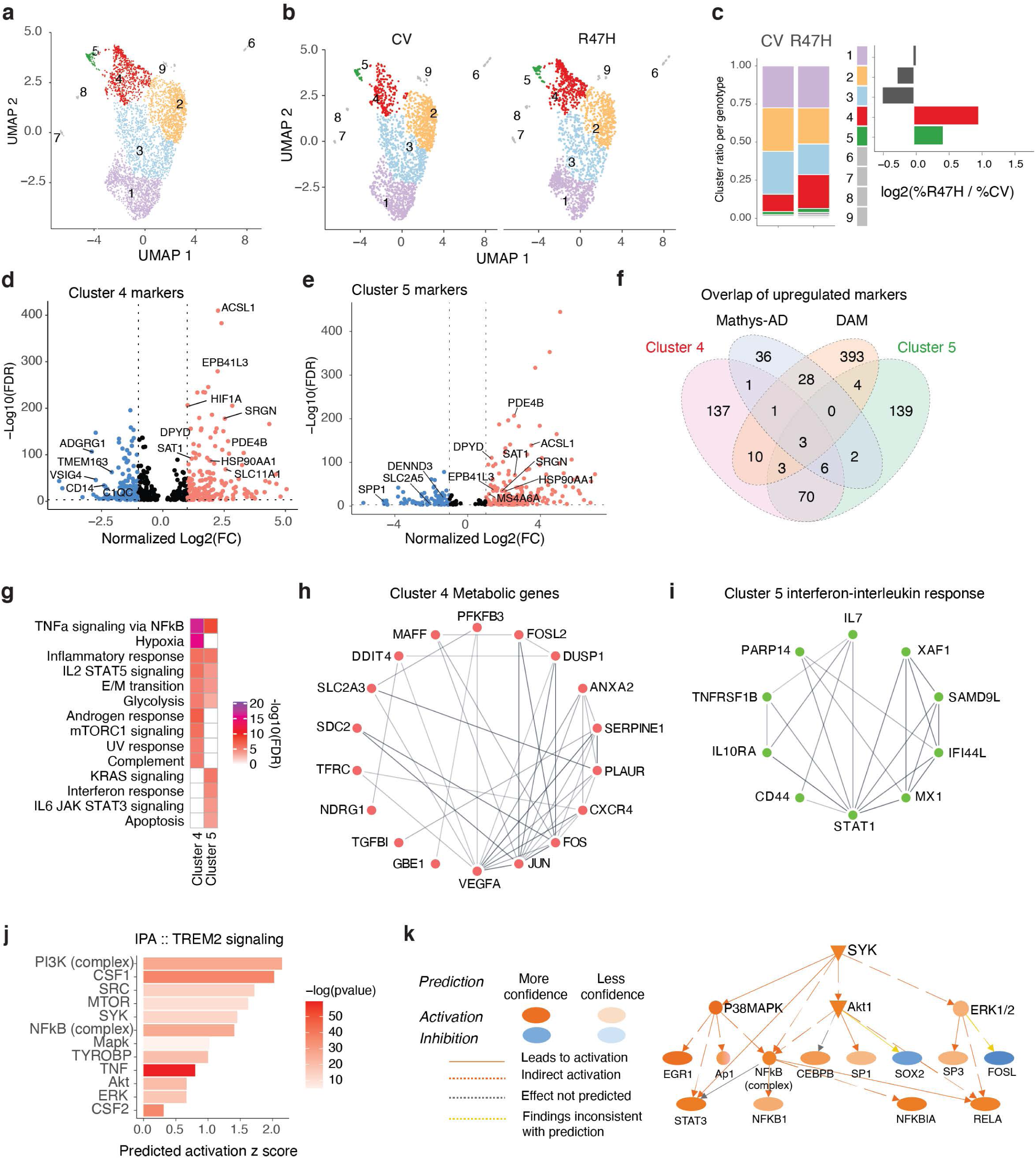
R47H Mutation Increases TREM2-Signaling in a Unique Microglia Cluster in Human AD Patient Brains. **(a-b)** UMAP of microglia subclusters for all nuclei (a) or split by TREM2 genotype (b). **(c)** Proportion of microglia subcluster per genotype (left) and log2 ratio of fraction in R47H versus CV (right). **(d-e)** Volcano plot of significant DEGs (FDR < 0.05) between homeostatic cluster 1 compared to cluster 4 (d) or cluster 5 (e). Genes overlapping with Mathys et al. highlighted ^41^. See also Supplementary Table 4. **(f)** Venn diagram showing number of overlapping upregulated markers from (d) and (e), DAM signature ^34^ and Mathys AD-microglia signature ^41^. **(g)** Heatmap of GSEA Hallmark pathways based on the unique, non-overlapping markers for clusters 4 and 5 identified in (f). Colors denote -log10(FDR). **(h-i)** Protein-protein interaction networks determined through the STRING database for genes overlapping in metabolic pathways: Hypoxia, mTORC1, and glycolysis for cluster 4 pathways (h) and interferon-interleukin pathways: Interferon response, IL2, and IL6 signaling for cluster 5 pathways (i) based on (g). **(j)** IPA of genes involved in TREM2 signaling based on DEGs identified in (d) and (e). **(k)** Diagram of SYK-AKT activation networks predicted by IPA upstream regulator analysis in (j). See also Extended Data Fig. 2-3 and Supplementary Table 4.

We then compared human R47H microglia in AD brains (cluster 4 or 5) with previously-identified mouse DAM microglia ^34^, and identified some overlap as well as significant differences (DAM, Fig. 2f, Extended Data Fig. 3b). Notably, genes involved in microglial homeostasis (*CX3CR1, TMEM119, TGFBR1, CSF1R*), complement signaling (*C1QA*, *C1QB*, *C1QC*), and purinergic signaling (*P2RY12, P2RY13*, *GPR34,* and *ENTPD1*) were downregulated in human *TREM2^R47H^* microglia and mouse DAMs. Upregulated genes overlapping with DAM signatures included *NAMPT*, *ATF3*, *SRGN*, *ATP1B3*, *ENPP1, CD83*, and *LPL*. However, many genes were differentially-regulated in human *TREM2^R47H^* microglia and mouse DAM. Several genes upregulated in *TREM2^R47H^* microglia were downregulated in mouse DAMs (*ABCA9*, *KLF2*, S*TAB, MTSS1, GLUL, TNFRSF1B, ABCC3*), while a small subset of genes downregulated in *TREM2^R47H^* microglia, including *SPP1,* involved in injury repair, and *SOCS6*, a negative regulator of the Jak-Stat pathway, were upregulated in DAMs (Extended Data Fig. 3b).

The R47H-unique signatures not overlapping with previously-identified human AD-microglial signatures or mouse DAMs were further analyzed using pathway enrichment analysis (Fig. 2f). The top pathway in both cluster 4 and 5 was TNF-*α* signaling via NF-*κ*B, suggesting an elevated proinflammatory state (Fig. 2g). R47H-induced changes were also associated with metabolic pathways, such as hypoxia, glycolysis and mTORC1 signaling, and immune pathways, such as interleukin (IL2 and IL6) and interferon response (Fig. 2g), with many of these pathway genes being interconnected (Fig. 2h,i). Interestingly, upstream and downstream mediators of TREM2 signaling, including AKT and SYK, were predicted to be activated in these human R47H-enriched microglia (Fig. 2j,k). Together, in human AD patients, the R47H mutation expanded a unique microglial subpopulation characterized by elevated pro-inflammatory transcripts and reduced homeostatic microglial genes, and hyperactivation of TREM2- associated signaling molecules.

### R47H-*hTREM2* Exacerbates Spatial Learning and Memory Deficits in Female Tauopathy Mice

To further dissect the molecular pathways induced by the R47H mutation, we generated knock-in mouse lines expressing one copy of CV- (*hTREM2^CV/+^*) or R47H-*hTREM2* (*hTREM2^R47H/+^*) cDNA at the *mTrem2* locus using CRISPR (Fig. 3a). PCR and Sanger sequencing confirmed the correct recombination and insertion of human TREM2-CV and TREM2-R47H cDNA at the *mTrem2* locus (Extended Data Fig. 4a,c-e). We did not detect any non-specific integration in the *hTREM2^R47H/+^* mouse line. However, a non-specific integration event occurred in *hTREM2^CV/+^* mice at an unknown mouse genomic region (Extended Data Fig. 4b,f-g). Nevertheless, *hTREM2^CV/+^* and *hTREM2^R47H/+^* mice had equivalent expression levels of *hTREM2* mRNA (Fig. 3b) and protein (Fig. 3c,d), and *hTREM2^R47H^* mice showed a dose-dependent reduction of *mTrem2* mRNA (Fig. 3e), indicating successful replacement of *mTrem2* by *hTREM2*. The matched expression levels of CV- and R47H-*hTREM2* mRNA and protein allowed us to assess the consequences of heterozygous R47H variant *in vivo*.

**Figure 3.**
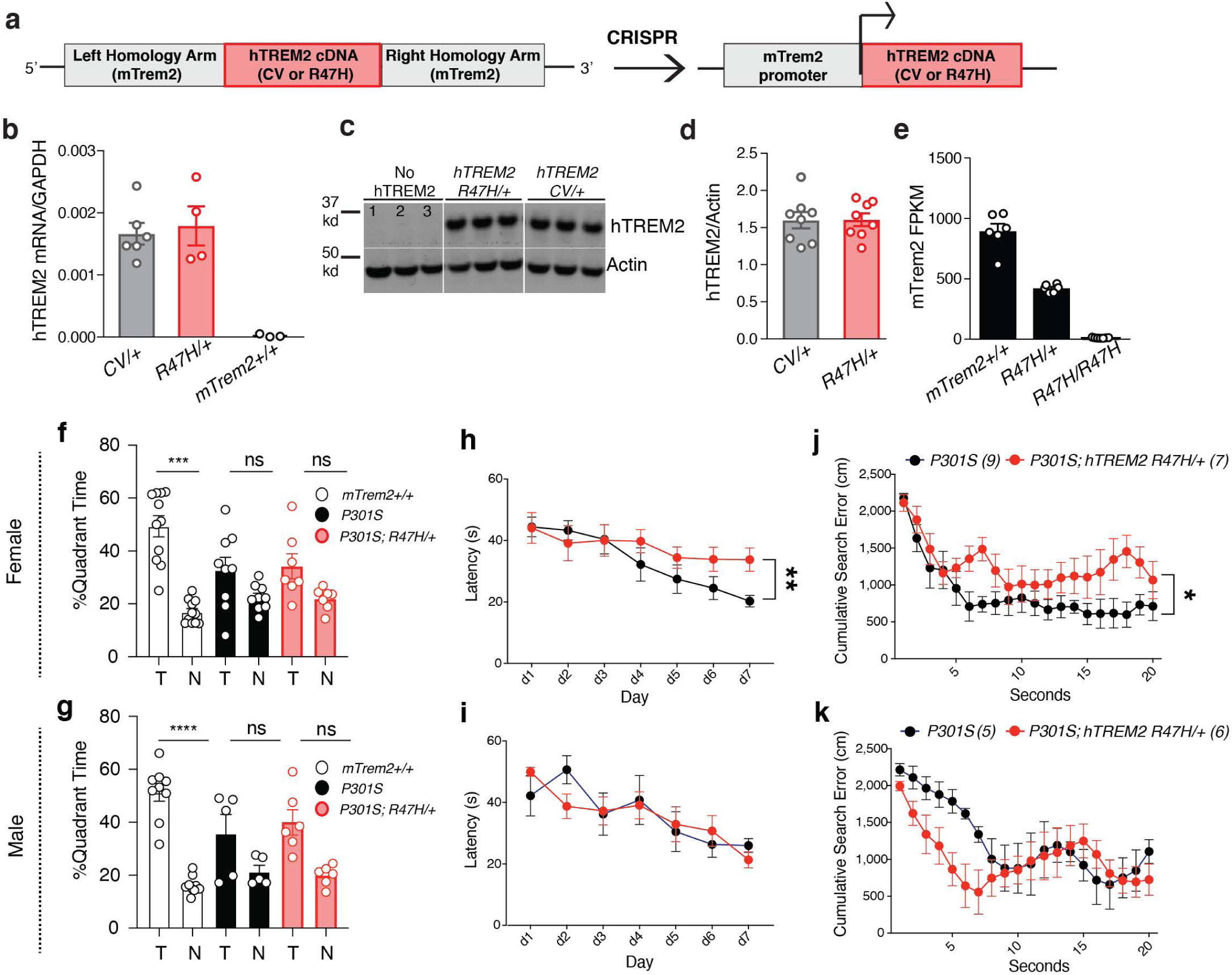
R47H-hTREM2 Exacerbates Spatial Learning and Memory Deficits in Female Tauopathy Mice. **(a)** The human *TREM2* donor vector was designed with two 1-kilobase long arms homologous to mouse *Trem2* (*mTrem2*) flanking the common variant (CV) or R47H human *TREM2* (*hTREM2)* cDNA sequence. When inserted into the genome, *hTREM2* cDNA is driven by the endogenous *mTrem2* promoter. **(b)** Quantitative real-time PCR analysis of cortical tissue from 3-4-month-old mice for *hTREM2* mRNA. Samples were run in triplicate, and averages of the three wells were used for quantification, normalized to *Gapdh*. Each dot represents the average of three wells from one mouse. Two-tailed Mann-Whitney U-test comparing CV/+ and R47H/+. **(c)** Representative western blot of RIPA-soluble cortical lysates from 8-9-month-old mice immunoblotted for hTREM2 and *β*-actin. Lane 1=*mTrem2*^-/-^, Lanes 2-3=*mTrem2*^+/+^. **(d)** Quantification of hTREM2 levels normalized by *β*-actin of the entire cohort by western blot. Student’s two-tailed t-test. **(e)** *mTrem2* mRNA expression level from RNA-seq of adult microglia isolated from whole brain of 3-4-month-old mice. FPKM, fragments per kilobase of transcript per million mapped reads. Kruskal-Wallis statistic=16.01, p=1.289e-007, Kruskal-Wallis test with Dunn’s post-hoc analysis. **(f,g)** Percentage of time spent in the target (T) or the average time spent in the nontarget (N) quadrants during the 24-hr probe for P301S *hTREM2^R47H/+^* females (**f**) and males (**g**) and their sex-matched *mTrem2^+/+^* and P301 *mTrem2^+/+^* littermate controls. ***t=6.088, p=0.0001 (f), ****t=7.834, p=5.077e-005 (g), paired two-tailed Student’s t-test for n>8, two-tailed Mann-Whitney U-test for n<8. **(h,i)** Latency to reach the platform during hidden trials (d1-d7) for female (**h**) and male (**i**) P301S *hTREM2^R47H/+^* and their P301S *mTrem2^+/+^* littermate control mice. **p=0.003, STATA mixed-effects modeling. **(j,k)** Cumulative search error for female (**j**) and male (**k**) P301S *hTREM2^R47H/+^* and their P301S littermate control mice. *U=9, p=0.0164, two-tailed Mann-Whitney U-test of area under the curve. Behavioral data represent the combination of two behavioral cohorts that were run independently. Values are mean ± SEM. See also Extended Data Fig. 4-7.

Tau pathology strongly correlates with cognitive deficits in AD ^53, 54^. To assess the effect of R47H- *hTREM2* on tau-induced cognitive deficits, we crossed *hTREM2^CV/+^* and *hTREM2^R47H/+^* mice with P301S mice, which express a human tau gene with the P301S mutation and develop hallmarks of tauopathy, including gliosis, tau inclusions, and cognitive deficits, such as deficits in hippocampal-dependent memory and spatial learning, mimicking human AD ^55^. The effects of a single allele of R47H-*hTREM2* on tau-induced spatial learning and memory were investigated using the Morris Water Maze. As expected, 7 to 9 month-old, non-transgenic mice preferred the target quadrant over the non-target quadrants while mice with P301S expression did not have a significant preference, indicative of spatial memory impairment (Fig. 3f,g). Interestingly, R47H-*hTREM2* significantly worsened spatial learning in female P301S mice, but had no statistically significant effect on males (Fig. 3h,i). P301S *hTREM2^R47H/+^* mice also made significantly more search errors during the 72-hour probe trial than controls (Fig. 3j), indicative of worsened spatial memory. In contrast, male P301S *hTREM2^R47H/+^* mice trended toward fewer search errors compared with male P301S mice during the probe trial (Fig. 3k). The spatial learning and memory curves of P301S *hTREM2^CV/+^* mice resembled that of P301S *mTrem2*^+/+^ (wild-type) mice regardless of sex (Extended Data Fig. 5a-d), suggesting the effects were R47H-mutation dependent. No differences were observed between genotypes in activity levels in the open field (Extended Data Fig. 5e-h) nor in the percentage of time spent in the open arms of the elevated plus maze (Extended Data Fig. 5i-l), ruling out hyperactivity and anxiety, which could confound the spatial memory test results.

Notably, the R47H mutation did not significantly impact the accumulation of insoluble tau aggregates detected using a conformation-specific antibody, MC1^56^, suggesting that the disease-enhancing effects of R47H-*hTREM2* in female P301S mice were not mediated by elevating toxic tau load (Extended Data Fig. 6). Indeed, even in the absence of tau pathology, the R47H mutation led to spatial learning deficits in females, without affecting baseline activity and anxiety levels (Extended Data Fig. 7). Taken together, our findings suggest that the R47H mutation renders hippocampal circuitry particularly vulnerable in both the absence and presence of tau pathology in a sex-dependent manner.

### R47H-*hTREM2* Increases Inflammation in Female Tauopathy Mice

To dissect how R47H*-hTREM2* microglia affects tau-mediated learning and memory, we next performed bulk-tissue RNA-seq of the hippocampus from 7- to 9-month-old male and female P301S *hTREM2^R47H/+^* and P301S *hTREM2^CV/+^* mice with matching *mTrem2* expression (Fig. 4a), and their littermate P301S *mTrem2*^+/+^ controls. Differential expression analysis revealed that R47H-*hTREM2* induced significant transcriptomic changes in females. 94 genes were upregulated by the mutation, including several DAM genes (e.g., *Ccl6, Clec7a, Siglec5, Cd9, Cd63*) ^34^ and other inflammatory genes (e.g., *Cxcl5, Ccl9*), while 28 genes were downregulated, including neuron-associated genes (e.g., *Adora2a, Syt6, Serpina9, Penk*) (Fig. 4b, Supplementary Table 5). In contrast, in male mice, R47H-*hTREM2* only significantly altered 3 genes in P301S mice (Fig. 4c), consistent with the lack of a significant effect of R47H-*hTREM2* on cognitive deficits in male P301S mice at this age (Fig. 3). The effects were specific to the R47H mutation, as CV-*hTREM2* had no significant transcriptional effect in female P301S mice and only altered 18 transcripts in male mice (Extended Data Fig. 8).

**Figure 4.**
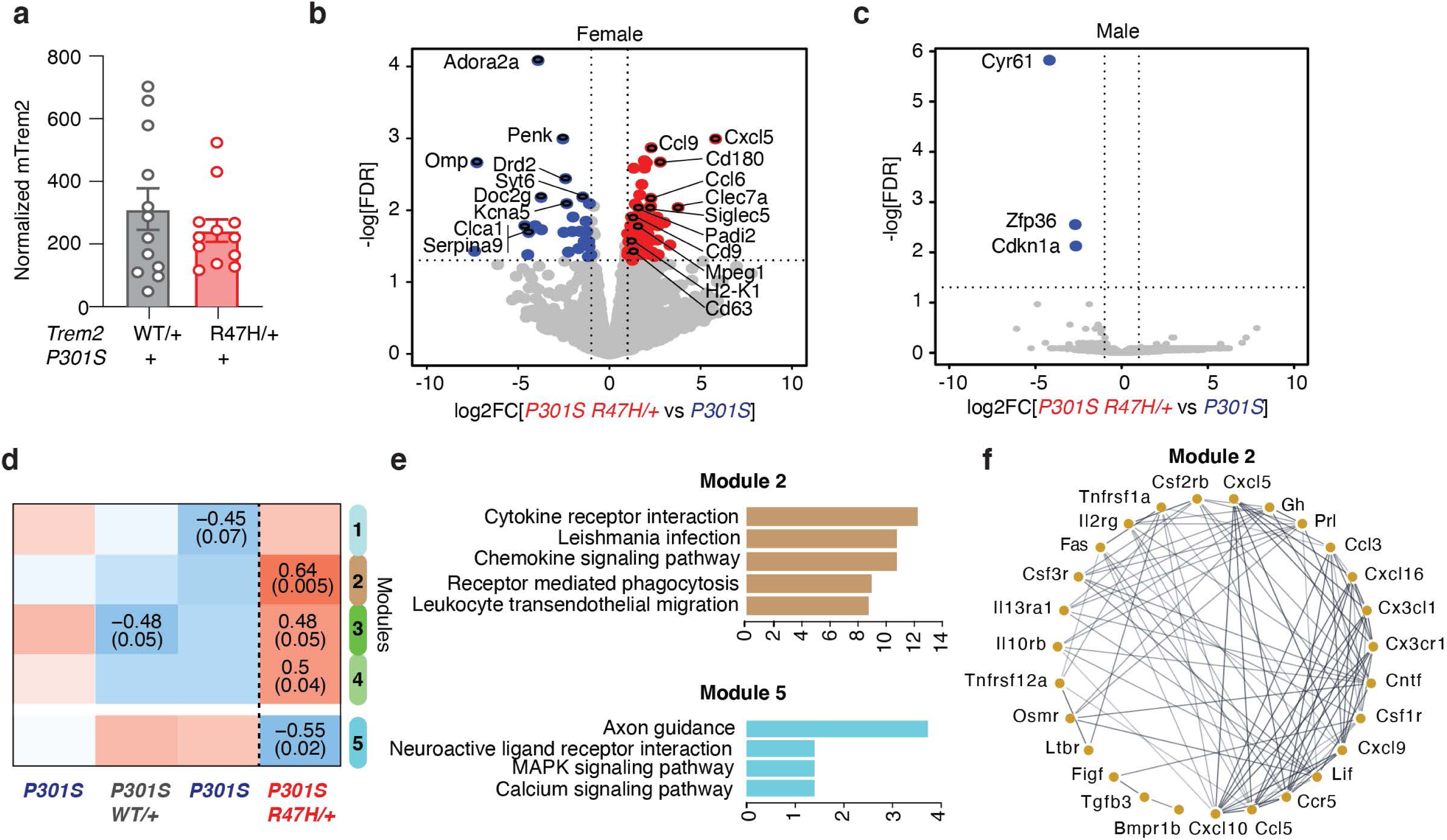
R47H-hTREM2 Increases Inflammatory Signatures in Female Tauopathy Mice. **(a)** Bar plot of normalized *mTrem2* RNA expression of bulk hippocampal tissue from 8–9-month-old P301S *hTREM2^R47H/+^* and P301S *hTREM2^CV/+^* mice. Student’s two-tailed t-test. **(b)** Volcano plot of bulk RNA-seq data of hippocampal tissue from 8–9-month-old female P301S *hTREM2^R47H/+^* mice and line-specific female P301S littermate controls. Blue dots are genes with significantly higher normalized counts in P301S controls than in P301S *hTREM2^R47H/+^* samples (28 mRNAs). Red dots are genes with significantly higher normalized counts in P301S *hTREM2^R47H/+^* samples than P301S controls (94 mRNAs). Highlighted upregulated genes are disease-associated microglial (DAM) genes and genes involved in inflammation while highlighted downregulated genes are the most significantly downregulated genes. Vertical dashed lines indicate log2FC ± 1. Horizontal dashed line indicates -log10(0.05). Wald test used. (n = 3 mice for P301S; n = 5 mice for P301S *hTREM2^R47H/+^*). See also Supplementary Table 5. **(c)** Volcano plot of bulk RNA-seq data of hippocampal tissue from male P301S *hTREM2^R47H/+^* and line-specific male P301S littermate controls. Blue dots are genes with significantly higher normalized counts in P301S controls than P301S *hTREM2^R47H/+^* samples (3 mRNAs). Vertical dashed lines indicate log2FC ± 1. Horizontal dashed line indicates -log10(0.05). Wald test used. (n = 2 mice for P301S; n = 5 mice for P301S *hTREM2^R47H/+^*). **(d)** Heatmap of results from weighted gene co-expression network analysis (WGCNA) of bulk RNA-seq data from (**b**) and Extended Data Fig. 8a, with only the significant module associations shown (top number: Pearson correlation, bottom number: adjusted p-value). Brown and cyan modules were the most significant. \*\**p* = 0.005, \**p* = 0.02. **(e)** Top 5 enriched KEGG (Kyoto Encyclopedia of Genes and Genomes) pathways of genes in brown and cyan modules from the WGCNA in (**d**). Colors of the bars represent the WGCNA module. See also Supplementary Table 6. **(f)** Network analysis using STRING database on genes from the KEGG “cytokine receptor interaction” gene set of the brown module. See also Extended Data Fig. 8 and Supplementary Table 5-6.

Using weighted gene-correlation network analysis (WGCNA), we examined the transcriptomes of female P301S *hTREM2^CV/+^* and P301S *hTREM2^R47H/+^* mice and their P301S *mTrem2^+/+^* littermate controls, and identified four statistically significant correlational modules related to P301S *hTREM2^R47H/+^* mice (Fig. 4d). Pathway analysis showed that the brown module, which exhibited the most significant positive correlation with the P301S *hTREM2^R47H/+^* genotype, was enriched with transcripts encoding cytokines/chemokines and cytokine receptors (e.g., *Ccr5, Ccl5, Ccl3, Cxcl5*) (Fig. 4e-f, Supplementary Table 6). The cyan module, which negatively correlated with the P301S *hTREM2^R47H/+^* genotype, was significantly enriched in transcripts encoding axon guidance molecules (e.g., *Sema6b, Sema3f, Epha8, Ephb6*) (Fig. 4e, Supplementary Table 6). Together, these data suggest an upregulation of pro-inflammatory transcripts and a concomitant decrease in neuronal signaling genes in female P301S *hTREM2^R47H/+^* mice compared to control mice.

### R47H-*hTREM2* Enhances Expression of Disease-associated Microglial Signature and Akt Signaling in Female Tauopathy Mice

We next performed single-cell RNA-seq using the Smart-Seq2 platform to specifically probe the effects of the R47H mutation on the microglial transcriptome ^57^. Microglia were isolated from the hippocampal tissue of 8-month-old female *mTrem2^+/+^*, *hTREM2^R47H/+^*, P301S *mTrem2^+/+^*, and P301S *hTREM2^R47H/+^* mice, gating on CD45^low^CD11b^+^ cells (Extended Data Fig. 9a-b). Out of the 1,480 cells that were sorted, 1,424 passed quality control (Extended Data Fig. 9c-g). *mTrem2* expression level was higher in *mTrem2^+/+^* microglia compared to the *hTREM2^R47H/+^* microglia, confirming the replacement of one allele of *mTrem2* (Extended Data Fig. 9h). Two distinct clusters were identified by unsupervised clustering of these 1,424 cells (Fig. 5a). While cluster 1 microglia were found in all 4 genotypes, cluster 2 microglia were mainly associated with the expression of P301S tau (Fig. 5b). Strikingly, *hTREM2^R47H^*^/+^ expression significantly increased the proportion of cluster 2 microglia in P301S tau mice (Fig. 5b-c). Compared to cluster 1 cells, cluster 2 cells have significant upregulation of transcripts including several DAM transcripts, such as *Clec7a, Ctsb, Axl, Cst7, Apoe,* and *Cd63* (Fig. 5d,e, Supplementary Table 7), consistent with the increased inflammation observed in the bulk-tissue RNA-seq data (Fig. 4b). While classical microglial genes, such as *Hexb*, were present in both clusters, the homeostatic microglial gene *P2ry12* was downregulated in cluster 2 cells (Fig. 5e). A direct comparison of cluster 2 marker genes versus DAM signature genes showed a statistically-significant positive correlation (Fig. 5f). Thus, in the presence of tau pathology, *hTREM2^R47H^*^/+^ enhances the DAM-like subpopulation and increases expression of *Trem2*-dependent microglial transcripts associated with neurodegeneration (MGnD), such as *Apoe, Itgax, Lpl, Axl,* and *Cst7* ^33, 34^ (red, Fig. 5f). We further examined the microglial *Apoe* expression in brain sections of P301S *hTREM2^R47H/+^* mice compared to P301S *hTREM2^+/+^* mice by RNAscope and confirmed that the proportion of microglia expressing *Apoe* was significantly increased in P301S *hTREM2^R47H/+^* mice (∼90%) compared to P301S *mTrem2^+/+^* (∼60%) in the dentate gyrus of the hippocampus (Fig. 5g-i).

**Figure 5.**
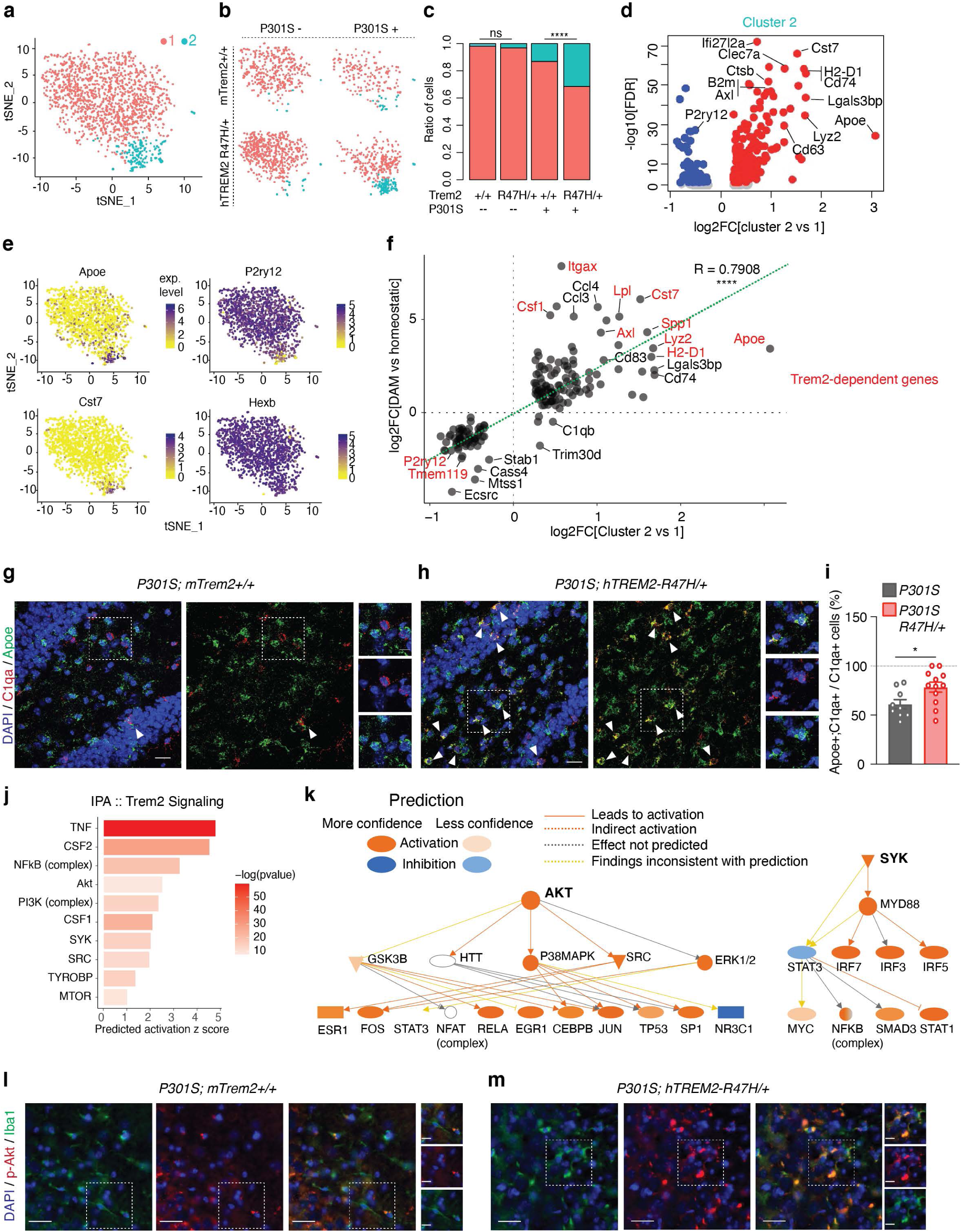
R47H-hTREM2 Enhances the Disease-Associated Microglia Population and Elevates Akt/Syk Signaling. **(a)** t-SNE plot of all 1,424 microglial cells analyzed and clustered. (n = 3 *mTrem2^+/+^*, 2 *hTREM2^R47H/+^,* 1 P301S, 2 P301S *hTREM2^R47H/+^*, 8-month-old female mice). **(b)** t-SNE plots based on clustering from (**a**) split by genotype. **(c)** Ratio of cells in each cluster by genotype. \*\*\*\**p* < 0.0001, two-sided Fisher’s exact test. **(d)** Volcano plot of DEGs defining cluster 2 compared to cluster 1. See also Supplementary Table 7. **(e)** Feature plots of transcript expression overlaid onto t-SNE of all microglial cells. Colored scale bar denotes normalized expression level. **(f)** Correlation scatterplot of DEGs in the microglial cluster 2 vs cluster 1 comparison (x-axis) compared to disease-associated microglia (DAM/MGnD) versus homeostatic microglia (y-axis) previously published ^34^. Red genes are *Trem2*-dependent. r = 0.7908, \*\*\*\**p* < 2.2e-16, Pearson’s correlation. **(g-h)** Representative images of RNAscope using probes against *C1qa* (red) and *Apoe* (green) of P301S **(g)** and P301S *hTREM2^R47H/+^* (**h**) dentate gyrus sections. White triangles highlight *C1qa*^+^;*Apoe*^+^ microglial cells. Dashed regions are zoomed in on the right side of the image. Scale bar = 20 μm, 10 μm for zoomed images. **(i)** Quantification of RNAscope images for percent of cells that are *C1qa*^+^;*Apoe*^+^ over total *C1qa*^+^ cells. *n* = 9 sections, 3 mice for P301S; 12 sections, 4 mice for P301S *hTREM2^R47H/+^*. Student’s t-test, \**p* = 0.0254, t = 2.426, df = 19. **(j)** Ingenuity Pathway Analysis (IPA) upstream regulator prediction for Trem2-signaling molecules based on cluster 2 markers from (d). Bar color denotes -log10(pvalue). **(k)** IPA example activated networks from (j) and their downstream predicted targets. **(l-m)** Representative images of immunostaining against phospho-Akt-Ser473 (red) and Iba1 (green) for P301S (**l)** and P301S *hTREM2^R47H/+^* (**m**) CA3 sections. Dashed regions are zoomed in on the right side of the image. Scale bar = 25 μm, 12.5 μm for zoomed images. Values are mean ± SEM. Each sequencing dataset represents one independent sequencing experiment. See also Extended Data Fig. 9-10 and Supplementary Table 7.

Upstream regulator analysis further predicted activation of Trem2-associated downstream signaling molecules, such as *Tyrobp, Akt*, *Syk* and *Src,* as well as activation of upstream regulators of Trem2 such as *Csf1* and *Csf2* ^7^ (Fig. 5j,k). Staining of brains from P301S *mTrem2^+/+^* and P301S *hTREM2^R47H/+^* mice against phospho-Akt-Ser473, the major phosphorylation site for Akt activation, showed colocalization of phospho-Akt signaling within Iba1+ microglial cells, with most of the phospho-Akt signal in P301S *mTrem2^+/+^* localized to non-Iba1+ cells (Fig. 5l). Consistent with the transcriptomic findings, an increased level of phospho-Akt in Iba1+ microglia was detected in P301S *hTREM2^R47H/+^* brains compared to P301S *mTrem2^+/+^* brains (Fig. 5l,m). Therefore, *hTREM2^R47H/+^* expression in female tauopathy mice induced similar features observed in AD *TREM2^R47H^* human microglia (Fig. 2j,k), including an expanded DAM-like subpopulation with enhanced Syk-Akt signaling and increased expression of a subset of transcripts previously found to be *Trem2*-dependent.

Aside from modulating the microglial inflammatory response, Trem2 is also involved in other key microglial functions. To test these, we assessed the microglial response to injury and phagocytosis ^12, 16, 36, 58^. The R47H mutation, however, did not alter the microglial response to laser-induced injury compared to *hTREM2^CV/+^* or *mTrem2*^+/+^ controls (Extended Data Fig. 10a-c, Supplementary Movie 1). The effects of R47H on phagocytosis were assessed by acquiring time-course images of primary microglia incubated with pH-rodo-conjugated *E. coli* substrates. Consistent with a previous study in HEK293 cells^29^, we detected no difference in the dynamics of fluorescence intensity over time among these three genotypes (Extended Data Fig. 10d-e), suggesting no change of phagocytic activity of *E. coli* induced by R47H. Due to heterogeneity in the phagocytosis kinetics between cells, we further performed a more detailed single-cell image tracking analysis by clustering cells based on different fluorescence dynamics, which correlated well with the bulk fluorescence analysis (Extended Data Fig. 10f). Again, *hTREM2^R47H/R47H^* microglia exhibited similar patterns of phagocytic behaviors at the single-cell level as *hTREM2^CV/CV^* and *mTrem2^+/+^* microglia (Extended Data Fig. 10g).

### Akt Activation Underlies Tau-mediated Proinflammatory Signatures in R47H-*hTREM2* Microglia

Our results so far showed that the R47H mutation enhances proinflammatory microglial responses in human AD and in female mouse tauopathy brains. We next investigated how the R47H mutation affects microglial response to tau specifically *in vitro*. To maximize the R47H effects, we treated *hTREM2^R47H/R47H^* microglia with tau fibrils and used *mTrem2*^+/+^ microglia as controls. Compared to *mTrem2*^+/+^ microglia, tau fibril stimulation significantly upregulated genes enriched in several signaling pathways in *hTREM2^R47H/R47H^* microglia (Fig. 6a,b). The cytokine–cytokine receptor interaction pathway was one of the top pathways altered by *hTREM2^R47H/R47H^* (Fig. 6b), consistent with our observation in female P301S *hTREM2^R47H/+^* mice (Fig. 4e). Moreover, Akt and Syk involved in Trem2-signaling pathways were again predicted to be activated in these microglia (Fig. 6c-e), similar to observations in our tauopathy mouse model and human AD tissues.

**Figure 6.**
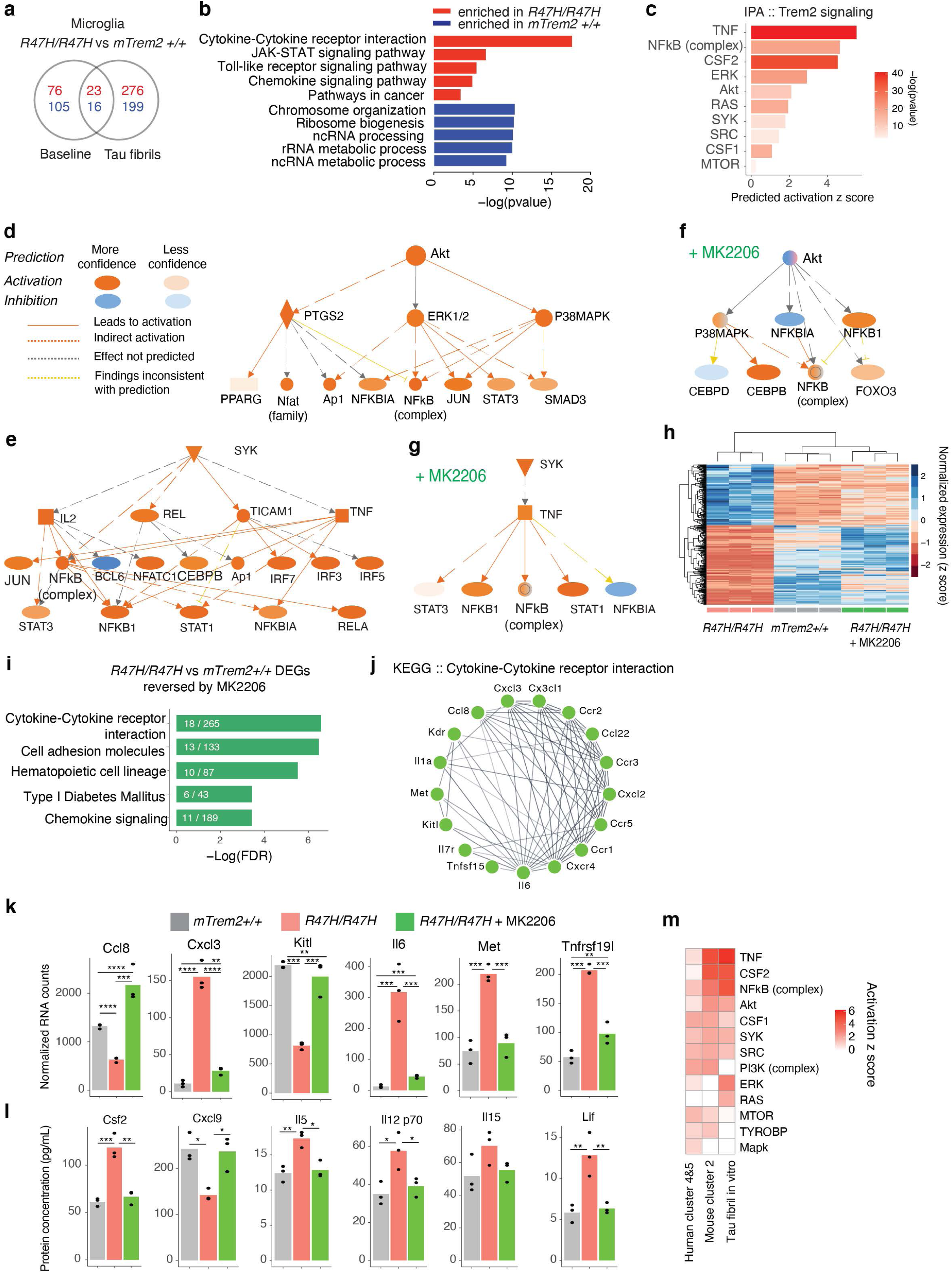
R47H-hTREM2 Primary Microglia Induce an Akt-dependent Pro-inflammatory Signature in Response to Tau Fibrils. **(a)** Venn diagram of differentially expressed genes between *mTrem2^+/+^* and *hTREM2*^R47H/R47H^ primary microglia with or without tau fibril stimulation. Red and blue numbers denote upregulated and downregulated genes, respectively. (n = 3 biological replicates for all conditions). **(b)** KEGG pathway enrichment analysis of the genes from (a) that were uniquely changed in *hTREM2^R47H/R47H^* microglia under tau fibril stimulation conditions. **(c)** IPA upstream regulator prediction for Trem2-signaling molecules based on genes that were uniquely changed in *hTREM2^R47H^*^/*R47H*^ microglia in response to tau fibril stimulation from (a). **(d-e)** IPA Akt (d) and Syk (e) activated networks from (c) and their downstream predicted targets. **(f-g)** IPA Akt (f) and Syk (g) networks and their predicted inhibition/activation states after MK2206 (Akt-inhibitor) treatment of *hTREM2^R47H^*^/*R47H*^ vs. *mTrem2^+/+^* microglia with tau fibril stimulation. **(h)** Heatmap showing z scores of normalized expressions of 477 genes that are changed by *hTREM2^R47H^*^/*R47H*^ compared to *mTrem2^+/+^* and are reversed towards control expression levels with MK2206 treatment. **(i)** KEGG pathway enrichment analysis of genes highlighted in (h). **(j)** STRING network representation of the genes in the “Cytokine-Cytokine receptor interaction” pathway from (i). **(k)** Barplots of example genes changed by *hTREM2^R47H^*^/*R47H*^ but reversed back to normal expression levels by MK2206. **(l)** Barplots of example cytokines measured by MAGPIX changed by *hTREM2^R47H^*^/*R47H*^ but reversed back to normal protein expression levels by MK2206. **(m)** Heatmap comparing the IPA predicted activation z score of TREM2 signaling molecules for all three models (Fig. 2, Fig. 5). See also Supplementary Tables 8-9.

Next, we directly tested the extent to which Akt signaling contributes to exaggerated inflammatory responses in tau-treated *hTREM2^R47H/R47H^* microglia. We acutely inhibited Akt in *hTREM2^R47H/R47H^* microglia cultures with MK-2206, an allosteric Akt-specific inhibitor ^59^ before treating with tau fibrils. Transcriptomic analysis showed that MK-2206 specifically inhibited the Akt pathway (Fig. 6f) without altering the Syk pathway (Fig. 6g). Strikingly, transcriptomic analysis demonstrated that, out of 838 DEGs (adjusted p value < 0.05, log2FC > 1 or < −1) between *hTREM2^R47H/R47H^* and *mTrem2^+/+^* microglia treated with tau fibrils, 477 of them were reversed towards *mTrem2*^+/+^ control levels upon Akt-inhibition (green columns, Fig. 6h, Supplementary Tables 8-9). These genes were enriched in pathways related to cytokine–cytokine receptor interaction (Fig. 6i-k). These transcriptional changes were further confirmed by measuring secreted cytokines in response to tau fibrils with a multiplex immunoassay. Out of the 26 cytokines altered by the R47H mutation, 11 of them were significantly rescued by MK-2206 (Fig. 6l). These results suggest that at both the RNA and protein levels, hyperactivation of Akt signaling may mediate some of the R47H-induced pro-inflammatory signatures in response to tau pathology. Remarkably, the predicted activation of TREM2-associated signaling pathways was consistent in all three R47H models – human AD carriers, mouse tauopathy mice, and primary mouse microglia treated with tau fibrils (Fig. 6m), supporting hyperactive Akt as one of the consistent mechanisms underlying the disease-enhancing property of R47H mutation in mouse and human.

## DISCUSSION

Compelling human genetic studies strongly suggest that maladaptive innate immune responses are associated with elevated risk of developing late-onset AD. Recent single-cell transcriptomic findings suggest that a subpopulation of microglia is enriched in response to AD-related pathologies (DAM or MGnD) ^33, 34^. Still, among the DAM signature genes, the identity of those that are disease-enhancing microglial genes (DEMs), disease-mitigating microglial genes (DMMs), or mere bystanders remains elusive. As the strongest immune-specific risk gene, the R47H-*TREM2* variant provides a unique model to help define immune-mediated DEMs in human AD. Through single-nuclei transcriptomic analysis of mid-frontal cortical tissues from 16 AD human patients carrying the R47H or CV variant of *TREM2*, we uncovered a microglia subpopulation enriched in AD R47H-*TREM2* carriers. We further established the disease-enhancing property of R47H-enriched microglia in a newly-developed R47H-*hTREM2* knock-in tauopathy mouse model by combining functional behavior studies and single-cell microglial transcriptomic analysis. We showed that in the context of tauopathy, R47H-*TREM2* microglia are characterized by a heightened inflammatory state, partially mediated by Akt hyperactivation.

In previous studies in amyloid mouse models, the homozygotic R47H mutation was found to dampen the microglial response to amyloid pathology, and correlated with increased neurotoxicity ^40, 47^. Paradoxically, in a tauopathy mouse model, the homozygotic R47H mutation was shown to be neuroprotective against neurodegeneration ^48^. Our current study provides the first line of evidence that the heterozygotic R47H mutation is disease-enhancing in the presence of tau pathology. While the R47H mutation did not impact general microglial functions such as phagocytosis and response to acute injury, the mutation exacerbated tauopathy-induced spatial learning and memory deficits in female tauopathy mice without affecting other cognitive domains, such as activity or anxiety levels. Importantly, this enhanced toxicity in our tauopathy model was not due to differences in tau-pathology load. Instead, transcriptomic profiling revealed that the toxic effects of R47H-*hTREM2* on tau-mediated cognitive deficits in female mice were associated with significant and profound transcriptional changes, particularly increased expression of pro-inflammatory genes. Meanwhile, the lack of toxic cognitive effects of R47H in male tauopathy mice was associated with few transcriptional changes. These findings support the notion that R47H-induced cognitive-deficits are driven by disease-enhancing microglial responses to stimuli including pathogenic tau, but not by directly influencing tau load itself.

Our study is the first to characterize the R47H-enriched microglial subpopulation in human AD brains and tauopathy mouse brains using single-cell or single-nuclei transcriptomic analyses. To identify specific microglial signatures induced by the R47H mutation in human brains, we ensured that samples were matched in AD pathology levels, *APOE* genotype, and sex. We discovered that, at the single-nuclei level, there was an increase in a subpopulation of microglia in R47H-AD brains characterized by heightened TNF-*α* signaling via NF-*κ*B and interferon responses, altered metabolic pathways, and enhanced AKT signaling. Similar to a previous observation of reduced microglial activation signatures in AD R47H-*TREM2* carriers using bulk-tissue RNA-seq ^46^, we found decreased expression of *AIF1* and *IRF8* in the R47H-associated microglial clusters. However, we also found overwhelming increases in expression of other immune-related genes, including interferon and interleukin-related genes (*IL7, CD44, IFI44L, TNFRSF1B)* and DAM-related genes (*NAMPT*, *ATF3*, *SRGN*, *ATP1B3*, *ENPP1, CD83*, *LPL)* ^34^, as well as increases in AD-associated genes defined previously such as (*SRGN, HIF1A, MS4A6A, SAT1, CTSB)* ^60^, suggesting a need to use a diverse set of marker genes to define the activation state of microglia in the human AD brain.

We showed that R47H-*TREM2* in human AD and in tauopathy mice exhibited heightened proinflammatory states, diminished homeostatic signature, and enrichment of DAM signature genes. These findings were in contrast to findings in *Trem2*-deficient microglia in amyloid and tau mouse models, which exhibit microglia in homeostatic states with blocked induction of DAM/MGnD signatures ^33, 34^. Our findings are also distinct from studies using homozygous R47H-*hTREM2* in 5XFAD mice ^47^ and in P301S mice ^48^, and provide compelling evidence that the heterozygotic R47H mutation does not phenocopy *TREM2* deficiency in humans, which results in Nasu-Hakola disease, and in mouse tauopathy. This distinction is important, since the vast majority of R47H carriers in AD are heterozygotes, and we previously demonstrated that *Trem2* haploinsufficiency can have opposing effects on tau pathology and microglial activation compared with *mTrem2^−/−^* ^36^. How exactly the homozygous and heterozygous R47H mutation differ in their downstream signaling is unclear. It is conceivable that the interaction of R47H- and CV-TREM2 could alter ligand-binding properties, including conformation of the hybrid receptor and ligand-selectivity, leading to heightened pro-inflammatory signaling. Nevertheless, the R47H-microglia from our tauopathy mice shared similar features to the R47H-associated human microglia, with enhanced TREM2-Akt-cytokine signaling and pro-inflammatory signatures as well as sex-specific transcriptomic differences, illustrating the strong relevance of our R47H knock-in mouse model to the human condition.

AD is strongly modified by sex, with many studies showing sex-differences in the incidence, prevalence, pathological findings, and disease progression rates ^61–64^. The sex-specific effects of R47H- *hTREM2* uncovered in our behavioral tests and transcriptomic studies in both mouse and human may be mediated by the differences between male and female microglial transcriptomes ^65–67^. Indeed, the R47H mutation led to disease-enhancing effects only in female, not male mice of the same age group. The lack of detrimental effects of R47H on male tauopathy mice is consistent with a recent study in which only male mice were used ^48^. In human tissues, we also observed sex-specific microglial changes induced by the R47H mutation, though there were greater transcriptomic changes in male compared to female microglia. The distinct sex-specific microglial response induced by R47H in mice and humans could potentially be explained by differences in their cognitive status. In the human studies, male and female patients were matched in their Cognitive Dementia Ratings, with both sexes having robust transcriptional changes induced by R47H-*TREM2* microglia, while in mice, the lack of transcriptional changes in microglia was correlated with male R47H tauopathy mice that exhibited little tau-mediated cognitive impairment.

In a previous study, induction of the MGnD-state by Trem2, including upregulation of *Apoe,* has been shown to be sex-specific ^33^. Interestingly, ApoE4 increases risk for late-onset sporadic AD to a greater extent in women ^68, 69^, and female ApoE4 knock-in mice have spatial memory deficits not seen in males ^70^. Microglia-derived ApoE is a major source of plaque-associated ApoE and is thought to be the driver of neurodegeneration in tauopathy mouse models ^71, 72^, suggesting that sex-specific differences in microglia may impact the sex-dependent effect of ApoE4 in AD pathogenesis ^73^. However, how R47H-*TREM2* and different *APOE* genotypes might interact to affect microglial function is unknown. We focused our current assessment of R47H effects in the same *APOE* background (*APOE3/4)* to control for potential influences of different *APOE* isoforms. Further studies are needed to confirm and extend our observations and to investigate the effects of different *APOE* isoforms.

Since DAMs exhibit responses that could be either protective or detrimental, blocking or enhancing all DAMs is unlikely to result in neuroprotection in AD patients. By identify disease-enhancing properties associated with the R47H mutation, strategies could be developed to suppress DEMs specifically and reprogram microglial responses more precisely. In our current study, *hTREM2^R47H/R47H^* microglia treated with tau fibrils also exhibited enhanced pro-inflammatory gene expression and hyperactive Akt-signaling, as observed in R47H microglia in human AD patients. Pharmacological inhibition of Akt corrected more than half of the genes altered by R47H, including cytokines, suggesting that Akt inhibitors could be explored as therapeutic strategies to reprogram disease-enhancing microglial states. While further *in vivo* studies need to be done in order to validate these findings, our study opens new avenues for developing microglia-targeted therapies for AD.

## ACKNOWLEDGEMENTS

We thank Dr. Junli Zhang from the Gladstone Institutes Transgenic Core for microinjections into embryos to generate knock-in human TREM2 mice; Allan Villanueva and Elizabeth Fan for assistance with genotyping the mice; Dr. Li-Huei Tsai and Dr. Yueming Li for their time discussing the data and feedback on the manuscript; Dr. Rosa Rademakers, and Dr. Nilufer Taner for genotyping and Christopher Fernandez De Castro for assistance with the human brain tissues; Rose Horowitz for help with immunostaining; and Stephen Ordway and Kathryn Claiborn for manuscript editing. All *in vivo* imaging was done by L.K., with helpful advice from Dr. Mario Merlini and Dr. Katerina Akassoglou, at the Gladstone Center for In Vivo Imaging Research. All immunohistochemistry imaging was done by F.A.S. and L.K. using the Gladstone Histology Core microscopes. Behavioral assays were done by F.A.S. and D.L. using equipment from the Gladstone Behavioral Core. RNA-sequencing was done by the UCSF Center for Advanced Technology (bulk-seq), and Weill Cornell Medicine Genomics and Epigenomics Core Facility (single-cell and single-nuclei RNA-seq). The authors are also grateful to the Flow Cytometry Core and the Integrated Genomics Operation at MSKCC for their assistance. This work was supported by the National Institute of Health Grants R01AG051390, U54NS100717, R01AG054214, and Rainwater Foundation (to L.G), the National Institute of Aging Grant F31 AG058505 (to F.A.S.), the National Institute of Aging Grant F30 AG062043-02 and National Institute of Health Grant T32GM007618 (to L.K.), the Early Postdoctoral Mobility Fellowship from the Swiss National Science Foundation P2BSP3 151885 (to H.M.), National Institute of Aging Grant K01 AG057862 (to T.E.T), National Institute of Mental Health Grant R01 MH110504 (to G.Y.), National Institute of Aging Grant R01 AG057555 (to L.X.), UK Dementia Research Institute, which receives its funding from DRI Ltd, funded by the UK Medical Research Council, Alzheimer’s Society and Alzheimer’s Research UK, Medical Research Council MR/N026004/1, Wellcome Trust 202903/Z/16/Z, Dolby Family Fund, National Institute for Health Research University College London Hospitals Biomedical Research Centre, Research Centre at University College London Hospitals NHS Foundation Trust and University College London (to J.H.), the National Institute of Aging Grants U01AG046170, RF1AG054014, RF1AG057440 and R01AG057907 (to B.Z.), and the Alan and Sandra Gerry Foundation (to L.M.).

## AUTHOR CONTRIBUTIONS

L.G. F.A.S, and L.K conceived and planned experiments. F.A.S., L.K. J.C.U., T.T., S.G., and L.G. designed experiments. F.A.S., L.K., J.C.U., L.F., D.L., G.K.C. H.M., X.J., Q.L., L.Z., S.G., W.L., M.Y.W., X.W., Y.Z., Y.Q.L., and T.T. performed experiments. L.K., H.M., Q.L., L.Z., X.N., F.G., M.T., Y.M.L., G.F., G.C., Z.C., G.Y., M.W., B.Z., L.X., M.B., L.M., and F.W. contributed experimental and analytical tools. F.A.S., L.K., J.C.U., L.F., D.L., T.T., H.M., F.G., M.T., G.C., Q.L., M.W., B.Z., M.B., G.K.C., X.W., Y.Q., S.G., and X.N. analyzed data. J.H. and D.W.D. provided human samples. Y.Z., D.L., M.Y.W., and Y.Q.L. helped maintain the mouse colony. L.K, F.A.S, and L.G. wrote the manuscript with input from all other authors.

## METHODS

### EXPERIMENTAL MODELS AND SUBJECT DETAILS

#### Mice

CRISPR/Cas9-mediated knock-in of the common variant (CV) or R47H human *TREM2* cDNA in place of *mTrem2* was done by injecting embryos with Cas9, short-guide RNA (sgRNA), and donor vectors (generated by PNA Bio). The human *TREM2* cDNA sequence was flanked on each side by 1-kb homology arms for the *mTrem2*. The sequences are as follows: *Trem2* targeted region 5′CCTGCTGCTGATCACAGGTGGGA and sgRNA sequence (antisense) 5′TCCCACCTGTGATCAGCAGCAGG. Potential off-target genes were identified with CRISPR off-target prediction software (http://www.crispor.tefor.net). There were no predicted off-targets for 1- or 2-basepair mismatches. CV hTREM2 and R47H hTREM2 lines were maintained independently and backcrossed to nontransgenic C57BL/6 mice for two to three generations, then crossed to *Cx3cr1*^GFP/GFP^ or P301S mice. *Cx3cr1*^GFP/GFP^ (https://www.jax.org/strain/005582) were crossed with CV or R47H hTREM2 knock-in lines to obtain *Cx3cr1*^GFP/+^*hTREM^R47H/+^*, *Cx3cr1*^GFP/+^*hTREM2^CV/+^*, and *Cx3cr1*^GFP/+^*mTrem2^+/+^* littermates for both lines. P301S transgenic mice (https://www.jax.org/strain/008169) were crossed with CV or R47H hTREM2 knock-in mice to generate P301S *hTREM2^R47H/+^* and littermate P301S *mTrem2^+/+^* mice, as well as P301S *hTREM2^CV/+^* and littermate P301S *mTrem2^+/+^* mice. Mice of both sexes were used, and analyses based on sex are included in the main and supplementary figures. Mice underwent behavioral testing at 7 to 9 months of age and had not been used for any other experiments. At 8 to 9 months of age, the same mice were used for pathology and RNA-seq studies. *Cx3cr1*^GFP/+^ mice were studied at 12 to 17 months of age. All mouse protocols were approved by the Institutional Animal Care and Use Committee, University of California, San Francisco and Weill Cornell Medicine.

#### Human Postmortem Samples

Tissues from the mid-frontal cortices from brains of donors with AD and the R47H mutation and donors with AD but without the R47H mutation (common variant; CV) (n = 8 donors per group) were used for single-nuclei RNA-sequencing, for a total of 16 samples. Samples were matched in age, tau and amyloid pathology burden, *APOE* genotype, and Clinical Dementia Ratings. Samples were obtained fro mteh Mayo Clinic brain bank and derived from several different studies: State of Florida Alzheimer’s Disease Initiative (ADI), Alzheimer’s cases derived from the Mayo Clinic (ADC), and cases obtained from outside sources, usually because of atypical clinical syndromes (such as corticobasal syndrome, frontal lobe dementia or progressive aphasia), but AD as the underlying pathology (Consult). Mayo Clinic brain bank operates under procedures approved by the Mayo Clinic institutional review board (IRB# is 15-009452). All brains were donated after consent from the next-of-kin or an individual with legal authority to grant such permission. The institutional review board has determined that clinicopathologic studies on de-identified postmortem tissue samples are exempt from human subject research according to Exemption 45 CFR 46.104(d)(2).

Additional information about the donors can be found in Supplementary Table 1.

#### Isolation of Nuclei from Frozen Postmortem Human Brain Tissue

The protocol for isolating nuclei from frozen postmortem brain tissue was adapted from a previous study with modifications ^41^. All procedures were done on ice or at 4°C. In brief, postmortem brain tissue was placed in 1500 µl of Sigma nuclei PURE lysis buffer (Sigma, NUC201-1KT) and homogenized with a Dounce tissue grinder (Sigma, D8938-1SET) with 20 strokes with pestle A and 15 strokes with pestle B. The homogenized tissue was filtered through a 35-µm cell strainer, and were centrifuged at 600 *g* for 5 min at 4°C and washed three times with 1 ml of PBS containing 1% BSA, 20 mM DTT and 0.2 U µl^−1^ recombinant RNase inhibitor. Then the nuclei were centrifuged at 600 *g* for 5 min at 4°C and re-suspended in 800 µl of PBS containing 0.04% BSA and 1x DAPI, followed by FACS sorting to remove cell debris. The FACS-sorted suspension of DAPI-stained nuclei were counted and diluted to a concentration of 1000 nuclei per microliter in PBS containing 0.04% BSA.

#### Droplet-based Single-nuclei RNA-seq of Human Brain Tissue

For droplet-based snRNA-seq, libraries were prepared with Chromium Single Cell 3’ Reagent Kits v3 (10x Genomics, PN-1000075) according to the manufacturer’s protocol. The snRNA-seq libraries were sequenced on the NovaSeq 6000 sequencer (Illumina) with 100 cycles.

#### Analysis of Droplet-Based Single-nuclei RNA-seq Data from Human Brain Tissue

Gene counts were obtained by aligning reads to the hg38 genome with Cell Ranger software (v.3.1.0) (10x Genomics). To account for unspliced nuclear transcripts, reads mapping to pre-mRNA were counted. Cell Ranger 3.1.0 default parameters were used to call cell barcodes. We further removed genes expressed in no more than 2 cells, cells with unique gene counts over 4,000 or less than 200, and cells with high fraction of mitochondrial reads (> 5%). Potential doublet cells were predicted using DoubletFinder ^49^ for each sample separately with high confidence doublets removed. Normalization and clustering were done with the Seurat package v3.0.1 ^74^. In brief, counts for all nuclei were scaled by the total library size multiplied by a scale factor (10,000), and transformed to log space. A set of 2000 highly variable genes were identified with *SCTransform* from *sctransform* R package in the variable stabilization mode. This returned a corrected unique molecular identifiers (UMI) count matrix, a log-transformed data matrix, and Pearson residuals from the regularized negative binomial regression model. Principal component analysis (PCA) was done on all genes, and *t*-SNE was run on the top 20 PCs. Cell clusters were identified with the Seurat functions FindNeighbors (using the top 20 PCs) and FindClusters (resolution = 0.02). In this analysis, the neighborhood size parameter pK was estimated using the mean-variance normalized bimodality coefficient (BCmvn) approach, with 20 PCs used and pN set as 0.25 by default. For each cluster, we assigned a cell-type label using statistical enrichment for sets of marker genes ^50, 51^ and manual evaluation of gene expression for small sets of known marker genes. Differential gene expression analysis was done using the FindMarkers function and MAST^75^. To identify gene ontology and pathways enriched in the differentially expressed genes (DEGs), we tested enrichment of gene annotation collections from the Molecular Signatures Database (MSigDB) gene annotation database v6.1 ^76, 77^. For convenience, the MSigDB gene set collections have been assembled into an R package called “msigdb” which is publicly available from https://github.com/mw201608/msigdb. Enrichment analysis P values were calculated using the hypergeometric test (equivalent to the Fisher’s exact test, FET) in R language. To control for multiple testing, we employed the Benjamini-Hochberg (BH) approach ^78^ to constrain the false discovery rate (FDR).

##### Microglia Subclustering

The subset() function from Seurat was used to subset microglia-only cells. Seurat object was converted to Monocle3 ^79–81^ cds for subclustering analysis as we found the integration between samples was most optimal using Monocle3. Cluster marker genes were determined using fit_models(model_formula_str = “∼cluster”).

#### Quantitative Reverse-Transcription PCR

Flash-frozen cortices were thawed and homogenized with a 21G needle in RLT buffer with 1% β-mercaptoethanol. RNA was isolated with the RNeasy Mini-Kit (Qiagen), and the remaining DNA was removed by incubation with RNase-free DNase. Purified mRNA was then converted to cDNA with the iScript cDNA Synthesis Kit (Bio Rad). Quantitative RT-PCR was performed on the ABI 7900 HT sequence detector (Applied Biosystems) with PowerUp SYBR Green master mix (ThermoFisher Scientific). The average value of three replicates for each sample was expressed as a threshold cycle (C_t_), the point at which the fluorescence signal starts to increase rapidly. Then, the difference (ΔC_t_) between C_t_ values for the transcript of interest and for mouse *Gapdh* was calculated for each sample. The relative gene expression for each sample was calculated as 2^−ΔCt^. The following primers were used:

Primer: Human TREM2 Fwd: CCGGCTGCTCATCTTACTCT
Primer: Human TREM2 Rev: GGAGTCATAGGGGCAAGACA
Primer: Mouse GAPDH Fwd: TGGCCTTCCGTGTTCCTAC
Primer: Mouse GAPDH Rev: GAGTTGCTGTTGAAGTCGCA

#### Western Blot

Mouse brains were homogenized in RIPA buffer containing 50 mM Tris, pH 7.5, 150 mM NaCl, 0.5% Nonidet P-40, 1 mM EDTA (ThermoFisher Scientific), 1 mM phenylmethyl sulfonyl fluoride, protease inhibitor cocktail (Millipore Sigma) and phosphatase inhibitor cocktail (Millipore Sigma). After sonication, brain lysates were centrifuged at 18,000 *g* at 4°C for 30 min. Supernatants were collected and protein concentrations were measured with the Pierce BCA Protein Assay Kit (ThermoFisher Scientific). The same amount of protein was loaded onto a 4–12% SDS-PAGE gel (Invitrogen), transferred to nitrocellulose membranes (GE Healthcare), blocked with 5% milk, and immunoblotted in 2% milk. Bands in immunoblots were visualized by enhanced chemiluminescence (ThermoFisher Scientific) and quantified by densitometry with ImageJ (NIH). The antibody used for western blot was anti-TREM2 (1:500, Cell Signaling). Immunoreactivity was detected with goat anti-rabbit HRP (1:2000, Millipore Sigma).

#### Isolation of Adult Microglia

Adult microglia were isolated from 3- to 4-month-old *mTrem2*^+/+^, *hTREM2^R47H/+^*, and *hTREM2^R47H/R47H^* mice as described ^36^. Briefly, after perfusion, brains were chopped with a razor blade, incubated with 3% collagenase type 3 (Worthington), 3 U/ml dispase (Worthington) and DNase (Millipore Sigma) at 37°C, inactivated with 2.5 mM EDTA (ThermoFisher Scientific) and 1% fetal bovine serum (FBS) (Invitrogen), filtered through a 70-µm filter, centrifuged at 300 *g* for 5 min at 18°C and resuspended in fluorescence-activated cell sorting (FACS) buffer. Samples were incubated with myelin-removal beads (Miltenyi Biotec) for 15 min at 4°C, passed through a magnetic LD column (Miltenyi Biotec), centrifuged at 300 *g* for 10 min, and resuspended in FACS buffer. Cells were magnetically separated and sorted with CD11b beads (Miltenyi Biotec) and a magnetic MS column (Miltenyi Biotec). CD11b-positive cells were centrifuged at 300 *g* for 10 min, and RNA was extracted for RNA-seq.

#### Behavioral Tests

In all behavioral tests, *hTREM2^CV/+^* and *hTREM2^R47H/+^* mice were compared to their respective nontransgenic or P301S littermates. Experimenters were blinded to mouse genotypes throughout the experiments. Male and female mice were tested on separated days.

##### Morris Water Maze

The water maze consists of a pool (122 cm in diameter) containing opaque water (20 ± 1°C) and a platform (10 cm in diameter) 1.5 cm below the surface. Three different images were posted on the walls of the room as spatial cues. Hidden platform training (days 1–7) consisted of 14 sessions (two per day, 2 hrs apart), each with two trials. The mouse was placed into the pool at alternating quadrants for each trial. A trial ended when the mouse located the platform or after 60 sec had elapsed. At 24 and 72 hrs after training, the mice were tested in probe trials, in which the hidden platform was removed and mice were allowed to swim for 60 sec. Mice received 7 days of hidden platform training before the 24-hr and 72-hr probe trials. Visible platform testing was done 24 hrs after the last probe trial. Performance was measured with an EthoVision video tracking system (Noldus Information Technology).

##### Elevated Plus Maze

The maze consists of two 15 x 2-inch open arms without walls and two closed arms with walls 6.5 inches tall and is 30.5 inches above the ground. Mice were moved to the testing room 1 hr before testing to acclimate to the dim lighting. Mice were individually placed in the maze at the intersection of the open and closed arms and allowed to explore the maze for 10 min.

##### Open Field

Mice were individually placed into brightly lit automated activity chambers equipped with rows of infrared photocells connected to a computer (San Diego Instruments). Open field activity was recorded for 5 min. Recorded beam breaks were used to calculate total time of activity.

#### Immunohistochemistry and Image Analysis

Mice were transcardially perfused with phosphate-buffered saline (PBS). The brains were cut vertically into hemibrains. Half of each brain was flash frozen at –80°C for RNA-seq analyses; the other half was placed first in 4% paraformaldehyde for 48 hr and then in a 30% sucrose solution for 48 hr at 4°C and cut into 30-µm-thick sections with a freezing microtome (Leica). For Extended Data Fig. 6, free-floating sections (8–10 per mouse) were washed in PBS, placed in sodium citrate buffer for 30 min at 90°C for antigen retrieval, permeabilized with PBS containing 0.5% Triton X-100 for 10 min, and blocked in PBS containing 10% normal goat serum for 1 hr. Sections were then placed in PBS with 5% normal goat serum and primary antibodies overnight at 4°C. The next day, sections were washed in PBS containing 0.1% Triton X-100, incubated with Alexa-conjugated secondary antibodies in PBS with 5% normal goat serum and washed in PBS containing 0.1% Triton X-100. Images were acquired with a Keyence BZ-X700 microscope and a 10x objective. Immunoreactivity was quantified with ImageJ software (NIH). Antibodies used for staining were anti-MC1 (1:500, kind gift from Dr. Peter Davies) and anti-Iba1 (1:500, Wako). Secondary antibodies used for staining were donkey anti-mouse 488 and donkey anti-rabbit 546 (both 1:500, ThermoFisher Scientific).

For Fig. 5, free floating sections were washed in Tris-buffered saline (TBS) buffer followed by incubation in Antigen decloaker reagent (Biocare Medical, Cat#RV1000M) in TBS at 95°C three times for 5 min each time, shaking at 600rpm. They there then washed with TBS containing 0.5% Triton x-100 (TBST), brain sections were incubated with 10% BSA in TBST for 1 hour at room temperature followed by incubation at 4°C overnight with anti-Iba1 antibody (goat, 1:500, Abcam #Ab5076) and anti-phospho-Akt-Ser473 antibody (rabbit, 1:100, Cell signaling #9271) in TBST containing 0.5% BSA. Next day, brain sections were washed with TBST three times and incubated for 1 hour at room temperature with donkey anti-goat secondary antibody conjugated with Alexa Fluor 488 (1:500, Invitrogen, #A11055) and donkey anti-rabbit secondary antibody conjugated with Alexa Fluor 568 (1:500, Invitrogen, #10042) in TBST containing 0.5% BSA. Brain sections were washed three times with TBST, mounted on slides in Vectashield mounting reagent containing DAPI, air dried, and sealed with nail polish. Stained sections were imaged by Keyence BZ-X710 All-in-One Fluorescence Microscope using the 40x objective.

#### RNA-Seq and Analysis of Adult Microglia

RNA from adult microglia was extracted from *mTrem2^+/+^* (n = 7), *hTREM2^R47H/+^* (n = 6), and *hTREM2^R47H/R47H^* (n = 7) mice with the Qiagen RNeasy Mini Kit. RNA concentration was determined with a NanoDrop, and RNA quality was measured with a Bioanalyzer and Agilent RNA Pico Chip. Samples with an RNA integrity number >7 were considered of good quality and used for subsequent steps. Libraries were then prepared with the QuantSeq 3′ mRNA-Seq Library Prep Kit FWD for Illumina. Library quality was assessed with a Bioanalyzer and the Agilent High Sensitive DNA Chip. Individual library concentrations were measured with the Qubit dsDNA HS Assay Kit and submitted for SE50 sequencing on an Illumina HiSeq 4000. Quality control was done on base qualities and nucleotide composition of sequences. Alignment to the GRCm38.84 *Mus musculus* (mm10) refSeq (refFlat) reference gene annotation was done with the STAR spliced read aligner and default parameters. Counts were normalized with the median of ratios method.

#### RNA-Seq and Analysis of Bulk Hippocampal Tissue

Hippocampal RNA was isolated with the Qiagen RNeasy Mini Kit. After quality control analysis with a Bioanalyzer, the RNA was sent to Novogene for library preparation and PE150 sequencing with an Illumina HiSeq 4000 instrument. 48 samples were sent in for sequencing: 6 samples each from male and female P301S CV and R47H hTREM2 knock-in mice, and 3 samples each for male and female P301S *mTrem2*^+/+^ littermates of the *hTREM2^CV/+^* line and male and female P301S *mTrem2*^+/+^ littermates of the *hTREM2^R47H/+^* line.

##### Quality Control

We used multiple clustering methods to examine the quality of replicates and to identify possible outlier samples for exclusion if necessary. For the hTREM2 R47H line (LG72), three samples were excluded from further analysis based on clustering. Clustering techniques were applied to variance stabilizing transformed expression values, fragments per kilobase of transcript per million mapped reads, and values of log counts per million. The Pearson correlation coefficient was first used as a distance metric between samples. Hierarchical cluster analysis was then applied to measure similarity between the Pearson correlations. The hierarchical clustering algorithm was an iterative process. Each iteration joined the two most similar clusters (based on Pearson correlation) and computed the distance between remaining clusters, continuing until there is just a single cluster. The distances between clusters were computed at each stage using the complete linkage clustering method (see manual for R hclust function).

##### Differential Expression Analysis

Differential gene expression was calculated with the R package DESeq2 ^82^. Counts were normalized with the median of ratios [Anders & Huber 2010]. Genes with <15 counts across all samples were excluded from analysis. The false discovery rate (FDR) was calculated with the Benjamini-Hochberg method ^78^.

##### Weighted Gene Co-expression Network Analysis

Weighted gene co-expression network analysis (WGCNA) was done on normalized expression data with the R package WGCNA v1.51^83^. The top 5000 most variable genes were used to create modules, and the soft-thresholding power parameter was set to 14. The minimum module size was 30 genes, and modules with a module eigengene dissimilarity below 0.2 were merged, creating 11 modules of 43–1361 genes each after removal of the module (gray) that contain genes that do not belong to any other module. KEGG and GO biological processes were used for pathway analysis as described above.

##### Pathway Enrichment Analysis

Gene network analyses of RNA-seq data were done with gene set enrichment analysis (GSEA) ^76^; cell types were defined by the top 100 genes expressed by each CNS cell type ^84^. Pathway analysis was done with Gene Ontology (GO) biological processes ^85, 86^ and the Kyoto Encyclopedia of Genes and Genomes (KEGG) ^87^.

##### Network Visualization

Networks were visualized with Cytoscape (version 3.6.1) ^88^, the STRING database ^89^, and perfuse force directed layout.

#### Mouse Hippocampus Single-cell RNA-seq

##### Preparation of samples

Brain tissue was prepared using a previously published protocol ^57^. Briefly, 8- month-old female mice were anesthetized with avertin and transcardially perfused with phosphate-buffered saline (PBS). The brain without the cerebellum was harvested and collected into cold media with 15mM HEPES, 0.5% glucose in 1X Hanks’ Balanced Salt Solution without phenol red (ThermoFisher Scientific 13185-052) on ice. The entire procedure was done on ice. The hippocampus was extracted and chopped into pieces using a razor blade then further homogenized using a 2ml douncer containing 2ml medium A with 80uL DNase (12500 units/mL) and 5ul recombinant RNase inhibitor (Takara Bio 2313B). Homogenized tissue was filtered through a 70um strainers to obtain a single cell suspension. Cells were washed with medium A and resuspended in 850uL MACS buffer with 1.8uL RNase inhibitor (sterile-filtered 0.5% BSA, 2mM EDTA in 1 X PBS). Cells were incubated with 100uL myelin removal beads (MACS Miltenyl Biotec) for 10 min, and loaded onto LD columns (Miltenyi Biotec). Cells were collected and washed for FACS staining.

##### Single Cell Sorting for Single-cell RNA-seq

Cells were blocked in 5uL mouse Fc block for 5 min on ice then incubated with primary antibodies for 10 min then washed with FACS buffer (sterile-filtered 1% FCS, 2mM EDTA, 25mM HEPES in 1XPBS). Cells were incubated with secondary antibodies for 10min then washed with FACS buffer. Cells were resuspended in 500uL FACS buffer with RNase inhibitor (Takara Bio 2313B, 1:500) and 0.5ul Propidium Iodide (ThermoFisher Scientific P3566, 1:1000) for single cell index sorting. Cell sorting/flow cytometry analysis was done on the cell sorter (BD InFlux) at the Stanford FACS Facility. The following gates were used for sorting microglia: (1) forward scatter-area (FSC-A)/side scatter-area (SSC-A) (2) Trigger Pulse Width/ FSC (3) Live-Dead negative using PI (4) CD45^low^CD11b^+^ and CD45^hi^CD11b^+^. Single cells were sorted into 96-well plates containing 4uL lysis buffer containing 4U Recombinant RNase Inhibitor (Takara Bio 2313B), 0.05% Triton X-100, 2.5mM dNTP mix (ThermoFisher Scientific R0192), 2.5uM Oligo-dT30VN (50- AAGCAGTGGTATCAACGCAGAGTACT30VN-30), and ERCC Spike-ins (ThermoFisher Scientific 4456740) diluted at 1:2.4×10^7^. Plates were vortexed, spun down and frozen on dry ice, and plates were stored at −80°C freezer. Antibodies used for FACS: rabbit anti-mouse Tmem119 (Abcam ab210405, ∼200ug/ul, 1:400 dilution), CD45-PE-Cy7 (ThermoFisher Scientific 25-0451-82, 1:300), CD11b-BV421 (BioLegend 101236, 1:300), goat anti-rabbit Alexa 488 (ThermoFisher Scientific 11034, 1:300).

##### Single-cell RNA-seq library preparation

Sequencing libraries were prepared following the Smart-seq2 published protocol^90^. Briefly, plates were thawed and incubated at 72°C for 3 min in order to anneal RNAs to the Oligo-dT30VN primer. The 6uL of the reverse transcription mixture was added to each well: 95U SMARTScribe Reverse Transcriptase (100U/µl, Clontech 639538), 10U RNase inhibitor (40U/µl), 1XFirst-Strand buffer, 5mM DTT (Promega P11871), 1M Betaine (Sigma-Aldrich B0300-5VL), 6mM MgCl2, 1µM TSO (Exiqon, Rnase free HPLC purified). RT was performed at 42°C for 90 min, followed by 70°C, 5 min. 15ul of PCR amplification mix containing the following reagents was added to each well: 1X KAPA HIFI Hotstart Master Mix (Kapa Biosciences KK2602), 0.1uM ISPCR Oligo (AAGCAGTGGTAT CAACGCAGAGT), 0.56U Lambda Exonuclease (5U/ul, New England BioLabs M0262S). cDNA was amplified using the following PCR program: (1) 37°C 30 min; (2) 95°C 3 min; (3) 23 cycles of 98°C 20 s, 67°C 15 s, 72°C 4 min; (4) 72°C 5 min. cDNA samples were purified using PCRClean DX beads (0.7:1 ratio, Aline C-1003-50), and resuspended in 20ul EB buffer. cDNA quality was examined with a Fragment Analyzer (AATI, High Sensitivity NGS Fragment Analysis Kit:1 bp - 6000 bp, Advanced Analytical DNF-474-1000). To make libraries, all samples were diluted down to 0.15ng/ul in 384-well plates using Mantis Liquid Handler (Formulatrix) and Mosquito X1 (TTP Labtech) with customized scripts. Nextera XT DNA Sample Prep Kit (Illumina FC-131-1096) was used at 1/10 of recommendation volume, with the help of a Mosquito HTS robot for liquid transfer. Tagmentation was done in 1.6ul (1.2ul Tagment enzyme mix, 0.4ul diluted cDNA) at 55°C, 10 min. 0.4ul Neutralization buffer was added to each well and incubated at room temperature for 5 min. 0.8ul Illumina Nextera XT 384 Indexes (0.4ul each, 5uM from 4 sets of 96 indexes) and 1.2ul PCR master mix were added to amplify whole transcriptomes using the following PCR program: (1) 72°C 3 min; (2) 95°C 30 s; (3) 10 cycles of 95°C 10 s, 55°C 30 s, 72°C 1 min; (4) 72°C 5 min. Libraries from one 384 plate were pooled into an Eppendorf tube and purified twice using PCRClean DX beads. Quality and concentrations of the final libraries were measured with Bioanalyzer and Qubit, respectively. Libraries were sequences on the Illumina HiSeq 4000 at the Weill Cornell Medicine Genomics and Epigenomics Core Facility.

##### Processing of Single-cell RNA-seq Raw Data

Prinseq ^91^ v0.20.4 was first used to filter sequencing reads shorter than 30 bp (-min_len 30), to trim the first 10 bp at the 50 end (-trim_left 10) of the reads, to trim reads with low quality from the 30 end (-trim_qual_right 25) and to remove low complexity reads (-lc_method entropy, -lc_threshold 65). Then, Trim Galore v0.4.3 (https://github.com/FelixKrueger/TrimGalore) was applied to trim the Nextera adapters (–stringency 1). The remaining reads were aligned to the mm10 genome by calling STAR ^92^ v2.5.3a with the following options: –outFilterType BySJout,–outFilterMultimapNmax 20,–alignSJoverhangMin 8,– alignSJDBoverhangMin 1,– outFilterMismatchNmax 999,–outFilterMismatchNoverLmax 0.04,– alignIntronMin 20,–alignIntronMax 1000000,– alignMatesGapMax 1000000,–outSAMstrandField intronMotif. Picard was then used to remove the duplicate reads (VALIDATION_ STRINGENCY = LENIENT, REMOVE_DUPLICATES = true). Finally, the aligned reads were converted to counts for each gene by using HTSeq (-m intersection-nonempty, -s no) ^93^.

##### Quality Control for Single-cell RNA-seq Data

We used the following criteria to filter out cells with low sequencing quality. The distribution of total reads (in logarithmic scale) was fitted by a truncated Cauchy distribution, and data points in two tails of the estimated distribution were considered as outliers and eliminated. Fitting and elimination were then applied to the remaining data. This process was run iteratively until the estimated distribution became stable. The threshold was set to the value where the cumulative distribution function of the estimated distribution reaches 0.05. Cells with small numbers of detected genes and poor correlation coefficients for ERCC (low sequencing accuracy) were dropped. 1424 cells were retained for downstream analysis after filtering from 1480 cells.

##### Clustering Analysis of Single-cell RNA-seq Data

The Seurat R package v3.0.1 was used to perform unsupervised clustering analysis on the filtered scRNA-seq data ^74, 94^. Gene counts were normalized to the total expression and log-transformed. Principal component analysis was performed on the scaled data using highly variable genes as input. The JackStrawPlot function was used to determine the statistically significant principal components. These principal components were used to compute the distance metric and generate cell clusters. Non-linear dimensional reduction (t-SNE) was used to visualize clustering results. Differentially expressed genes were found using the FindAllMarkers function that ran Wilcoxon rank sum tests.

##### Ingenuity Pathway Analysis

DEGs and their log2-fold change expression values were inputted into IPA [citation]. For Trem2-signaling pathway predictions, previously published molecules involved in Trem2-signaling were manually curated and highlighted through network construction.

#### RNAscope

Mouse brains were harvested and quickly frozen on dry ice followed by OCT embedment. Brains were cut into 20 µm sections and mounted onto superfrost plus slides. The RNAscope assay was carried out with RNAscope Fluorescent Multiplex Assay (ACD, 320513). Briefly, after dehydration with ethanol, sections were permeabilized with Protease IV. Then samples were hybridized with RNA probes C1qa (ACD, 441221-C2) for activated microglia and Apoe (ACD, 313271-C3) at 40℃ for 2 h. Signal was amplified according to the instructions. After counterstain with DAPI, sections were mounted by Prolong Gold (Thermo Fisher, P36930). Line-scanning confocal images of hippocampal dentate gyrus region were obtained with a confocal microscope (LSM880, Carl Zeiss Microscopy, Thornwood, NY) and a 40x objective (1-µm focal plane intervals, one field of view per brain section) within a day post-mounting of sections. Images were examined by maximum intensity Z-projection and analysis was done using ImageJ v2.0.0 (NIH). All analysis and imaging were done blinded to the genotype.

##### Isolation of Primary Microglia - *in vitro* Tau Fibril Stimulation Assay and MK-2206 Treatment

Cortices was harvested from postnatal day 3 pups. The meninges were removed, and the cortical tissue was finely chopped with a razor blade and digested in 0.25% trypsin with DNAse (Millipore Sigma) at 37°C for 25 min. Digestion was stopped with DMEM containing 10% FBS. The tissue was then triturated and spun at 200 *g* for 5 min, and the pellet was resuspended in DMEM and 10% FBS and plated in T75 flasks that had been pre-coated with poly-D-lysine (Millipore Sigma) and rinsed with water. Mixed cultures were maintained in flasks for 10–11 days. The flasks were then shaken for 2 hr, spun at 200 *g* for 15 min, and the cells were plated at a density of 150,000/well in a 24-well plate. After 24 hr, cells were treated or not with 1 ug of 0N4R tau fibril per well (from Dr. Jason Gestwicki) for approximately 16 hr before lysis for RNA isolation. For drug treatment, cells were treated with either 1 μm MK-2206 (MedChem Express) suspended in DMSO or no drug with DMSO as a control. 6 hours after drug treatment, cells were treated with 1 ug of 0N4R tau fibril per well. 24 hours later, conditioned media for the MagPix assay was collected from the wells, spun at 4°C for 10 minutes at 3000 rcf, and the pellet was discarded to remove debris. A 32-plex MagPix Mouse Cytokine Chemokine assay (Milliplex) was run on the conditioned media. On the same day as conditioned media collection, RNA was isolated using the Mini Kit (Zymo Research) and sent to Novagene for library preparation and sequencing.

#### RNA-Seq and Analysis of Primary Microglia

RNA was extracted from samples with the Zymo *Quick*-RNA MiniPrep kit. For tau fibril treatment experiment without drug treatment, RNA concentration was determined with a NanoDrop, and RNA quality was measured with a Bioanalyzer and Agilent RNA Pico Chip. Samples with an RNA integrity number >7 were considered of good quality and used for subsequent steps. Libraries were then prepared with the QuantSeq 3′ mRNA-Seq Library Prep Kit FWD for Illumina. Library quality was assessed with a Bioanalyzer and the Agilent High Sensitive DNA Chip. Individual library concentrations were measured with the Qubit dsDNA HS Assay Kit and submitted for SE50 sequencing on an Illumina HiSeq 4000. Quality control was done on base qualities and nucleotide composition of sequences. Alignment to the GRCm38.84 *Mus musculus* (mm10) refSeq (refFlat) reference gene annotation was done with the STAR spliced read aligner and default parameters. Differential gene expression analysis was done with the R package DESeq2. Counts were normalized with the median of ratios method. Genes with <15 counts across all samples were excluded from analysis. The FDR was calculated with the Benjamini-Hochberg method. Pathway analysis was done using Gene Set Enrichment Analysis referencing Gene Ontology Biological Processes dataset. Predicted upstream activators of the transcriptome were determined using Ingenuity Pathway Analysis software (Qiagen).

#### In Vivo Imaging

For intravital imaging with two-photon microscopy, thinned-skull windows were made in 12–17-month-old *Cx3cr1*^GFP/+^ *hTREM^R47H/+^*, *Cx3cr1*^GFP/+^ *hTREM2^CV/+^*, and *Cx3cr1*^GFP/+^ *mTrem2^+/+^* as described previously ^36^. Briefly, mice were anesthetized, the skull was exposed, and a small area over the cortex was thinned manually and with a high-speed drill (K.1070 High Speed Rotary Micromotor drill; Foredom). Mice were fixed onto a custom-made head plate and imaged with an Ultima IV multiphoton microscope (Bruker) equipped with MaiTai DeepSee-eHP lasers (Spectra Physics) tuned to 920 nm for imaging and InSight X3 lasers (Spectra Physics) tuned to 880 nm for ablation. Z-stacks of images were acquired every 3 min in 1-µm steps with a 25x water-immersion objective at 2.4 optical zoom. Extensions and retractions of processes during baseline recordings were manually traced with the mTrackJ plugin. The movement of microglial cells toward a laser ablation site was analyzed by normalizing the number of processes near the injury site at each time point to the overall microglial density at that time point.

#### Primary Microglia Phagocytosis Experiment

##### Experimental and Imaging Set-up

Primary microglia were prepared as described above. Microglia were seeded at 30,000 cells / well in a 96-well plate. After 24 hours, media was replaced with FluoroBrite DMEM media (ThermoFisher) with 10% FBS and 0.015 mg pH-rodo *E. coli* (ThermoFisher; P35366) was added to each well. 4 fields of view were taken for each well every 30 minutes using Incucyte (Essen BioScience). Background fluorescence was subtracted and population-level fluorescence information was determined through the Incucyte analysis software using Top-Hat and threshold based on negative control well with no pH-rodo *E. coli*.

##### Single-cell image tracking. Model Training

The single-cell analysis pipeline was adapted from previous work, using the Usiigaci framework ^95^. A custom FIJI/ImageJ macro was written to manually annotate phase contrast images, such that a unique RGB value corresponded to an individual cell mask^96^. A preprocessing step converted the 8-bit RGB image into a 16-bit grayscale mask image (8-bit would yield max of 255 unique cells per image), such that each unique RGB cell mask was a unique grayscale value. A Mask R-CNN model implemented in Tensorflow and Keras (open-source Matterport Inc. – MIT License), was trained to segment cells in phase contrast microscopy images ^97–100^. The model had a learning rate of 1×10-3 for training and used a Resnet-101 backbone ^95, 99^. The network heads were trained first for 50 epochs and then the full network was trained for another 100 epochs. All model training was performed using a GPU on Google Colab. The imgaug package was used for data augmentation during training, incorporating image rotations and flips, along with brightness augmentations and gaussian blur augmentations stochastically. Transfer learning was performed based on final weights from Usiigaci model. The average of two different models (epochs 130 and 134) were used for instance segmentation during this study. These models were chosen based on lowest validation loss during training. The model was trained on 20 images with ∼50 cells per image to assess accuracy of the model on data that were not used during model training.

##### Analysis of Phase Contrast Microscopy Images

Phase contrast and fluorescent image were exported as TIFF images from the Incucyte system. Top-hat background subtraction was performed on the fluorescence image to remove autofluorescence. Time-lapse images were registered using a manual drift correction FIJI plug-in ^101^. The output from the Mask R-CNN model was fed into a K-D tree-based cell tracking pipeline based on trackpy, from the Usiigaci pipeline. Once cells from each frame were annotated, a python script was used to measure fluorescence of each cell over the entire time-lapse. A threshold was manually set for each individual experiment. Average intensity measurements above the threshold were measured for each individual cell by taking the total intensity within a cell above the threshold divided by the area of the cell above the set threshold. Cells with more than 30 missing time points and cells with more than 5 missing time points within the first 10 frames were excluded from the analysis and a linear regression was used to impute missing data from the time-lapse. Cells that contained enough datapoints to be included in the cut-off but missing datapoints at the beginning and end of the time-lapse were filled with zeros.

##### Clustering of Single-Cell Fluorescence Data

Clustering algorithms were adapted from the Seurat R package^74^ for single-cell RNA sequencing analysis. Briefly, the fluorescence measurement for each cell at each time point was scaled such that the mean expression across cells is 0 and all time points were used to generate a PCA. For clustering and generation of the tSNE, FindNeighbors, FindClusters (resolution 0.15), and RunTSNE functions were used.

#### Quantification and Statistical Analysis

Data were analyzed with GraphPad Prism v.7 (GraphPad Software, San Diego, California USA, www.graphpad.com), STATA12 (StataCorp. 2011. Stata Statistical Software: Release 12. College Station, TX: StataCorp LP), or R^102^. A multilevel mixed-effects linear regression model fitted with STATA12 was used to analyze latency in the Morris water maze. R was used to calculate the area under the curve for cumulative search errors in the Morris water maze. Outliers were removed with Prism’s outlier analysis algorithm. All statistical details can be found in the figure and figure legends. P < 0.05 and FDR < 0.05 was considered statistically significant, unless otherwise noted. All values are expressed as mean ± SEM, unless otherwise noted. A subset of mice from the behavior cohort was randomly selected for single-nuclei RNA-seq and bulk RNA-seq studies. Data and visualizations were done using ggplot2 ^103, 104^.

### DATA AND CODE AVAILABILITY STATEMENT

Full western blots (Fig. 3) are available as a source data files and RNA-seq gene lists with statistics are available in the respective supplementary tables accompanying this article. All RNA-seq data was deposited in the Gene Expression Omnibus (GEO) under the following series accession numbers: bulk RNA-seq of mouse hippocampus and primary microglia; GSE136389, human single-nuclei; GSE154578, mouse single-cell; GSE140670. If you are a reviewer, please consult the journal editor for the access token. Any code used for data analysis can be made available upon request.

### MATERIALS AVAILABILITY STATEMENT

Further information and requests for resources and reagents should be directed to and will be fulfilled by the Lead Contact, Li Gan.

## EXTENDED DATA FIGURE LEGENDS

**Extended Data Figure 1.**
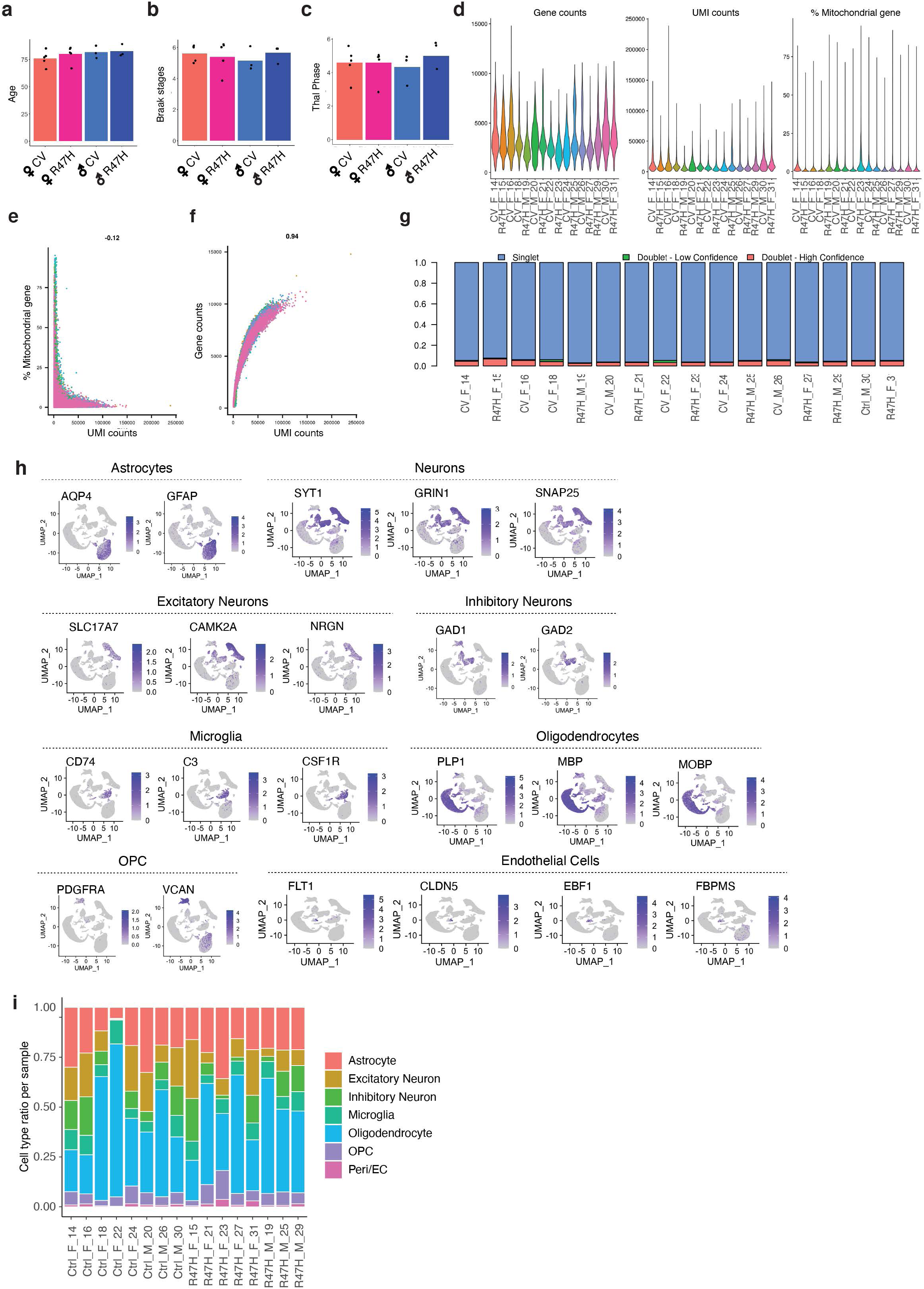
Quality Control Assessment of Single-Nuclei RNA-Seq of Human AD Patient Brain Tissues (Related to Fig. 1) **(a-c)** Bar plots of age (a), Braak stage (b), and Thal phase (c) of the patient tissues. **(d)** Violin plots showing spread of total genes, total UMIs, and percent of mitochondrial genes detected per nuclei for each individual sample. (**e-f**) Correlation between UMI counts and percentage of mitochondrial genes per nuclei (e) and total genes detected (f) for all samples. **(g)** Ratio of nuclei in each sample determined as singlets, doublets (low confidence), and doublets (high confidence). Only high-confidence doublets were removed for downstream analysis. **(h)** Feature plots showing expression of cell type marker genes for each cell type. **(i)** Proportion of different cell types captured per sample.

**Extended Data Figure 2.**
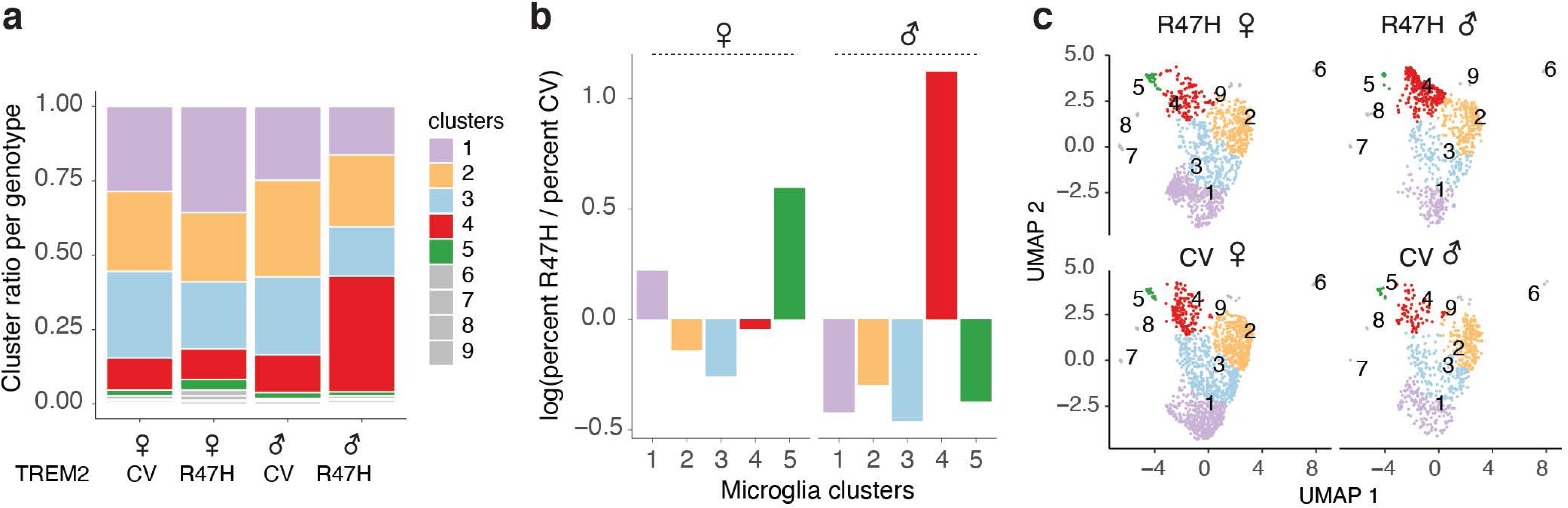
R47H-hTREM2 Induces a Sex-Specific Change in the Microglial Transcriptome of Human AD Patients (Related to Fig. 2). **(a)** Microglia subcluster ratio per genotype and sex. **(b)** Log2 ratio of microglia subcluster fraction in R47H versus CV for each sex. **(c)** UMAP split by genotype and sex. Colors denote subclusters from (a).

**Extended Data Figure 3.**
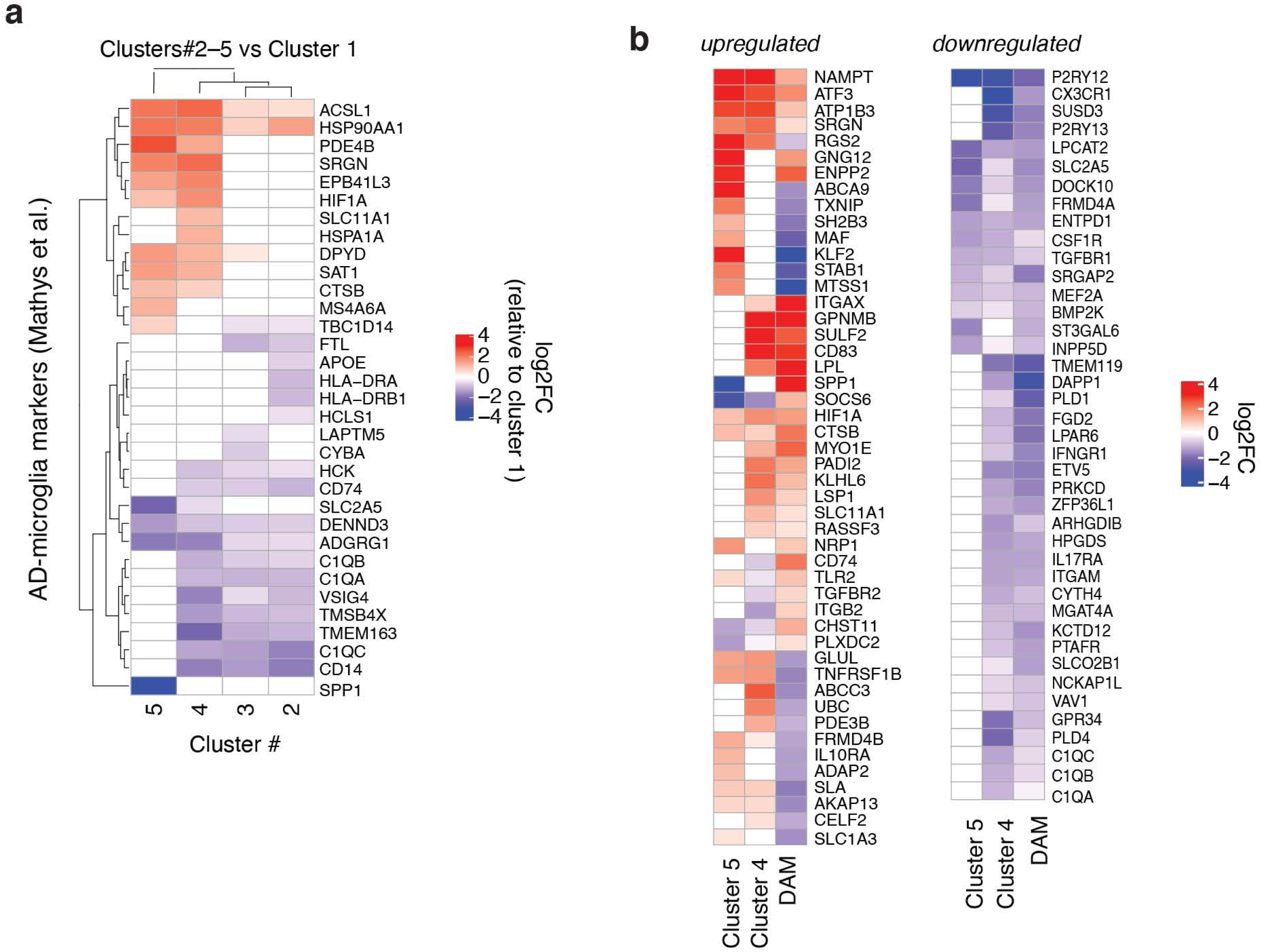
R47H-associated Microglia Signature Overlaps with Previously-identified AD-associated Signatures (Related to Fig. 2) **(a)** Heatmap showing Log2-fold change values of DEGs from (Fig. 2d) and (Fig. 2e) compared to AD-associated microglia signature identified in Mathys et al. (Mic1) ^41^. **(b)** Heatmap showing Log2-fold change values of DEGs from (Fig. 2d) and (Fig. 2e) compared to the DAM signature ^34^.

**Extended Data Figure 4.**
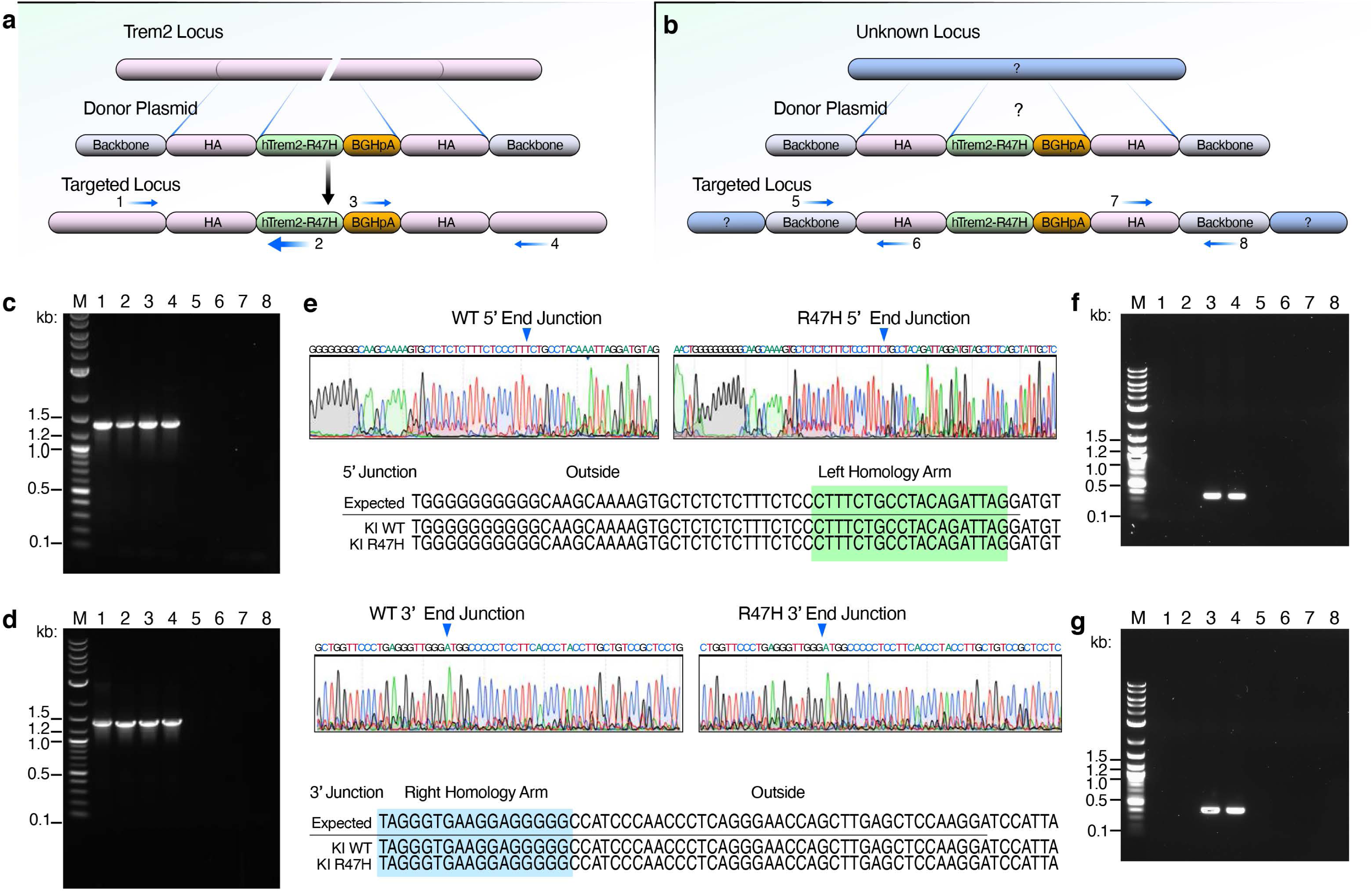
Characterization of CV-*hTREM2* and R47H-*hTREM2* Knock-in Mouse Lines (Related to Fig. 3) **(a)** Diagram of the strategy to detect the specific targeting of the R47H-*hTREM2* (or CV-*hTREM2*) into the mouse *Trem2* genomic locus and primer locations for detecting integration. PCR primers labeled as arrows and are located as follows: Primer 1 is located outside of the left homology arm, Primer 2 is on 5’ end of *hTREM2* cDNA, Primer 3 is on BGHpA, Primer 4 is located outside of the right homology arm. **(b)** Diagram of the strategy to detect non-specific integration of the R47H-*hTREM2* (or CV-*hTREM2*) cDNA in the mouse genome. PCR primers labeled as arrows and are located as follows: Primer 5 is on the vector, Primer 6 is on the left arm, Primer 7 is on the right arm, Primer 8 is on the vector. **(c)** PCR screening for the 5’ end specific homologous recombination of CV-*hTREM2* and R47H- *hTREM2* mice using Primers 1 and 2. Lane M = 1kb plus DNA ladder, Lanes 1-2 = R47H-*hTREM2*, Lanes 3-4 = CV-*hTREM2*, Lanes = 5-6 = C57BL/6 control, Lanes 7-8 = H_2_O control. The size of PCR product is about 1345 bp. **(d)** PCR screening for the 3’ end specific homologous recombination of CV-*hTREM2* and R47H- *hTREM2* mice using Primers 3 and 4. Lane M = 1kb plus DNA ladder, Lanes = 1-2 = R47H-*hTREM2*, Lanes 3-4 = CV-*hTREM2*, Lanes 5-6 = C57BL/6 control, Lanes 7-8 = H_2_O control. The size of PCR product is about 1339 bp. **(e)** Sanger sequencing chromatograms for the junctions. Sequences of the intended and obtained knock-in junctions in the mouse lines show correct homologous recombination. 5’ end of left homology arm is highlighted in green and the 3’end of right homology arm is highlighted in blue. **(f)** PCR detection of non-specific integration on 5’ end of the CV-*hTREM2* and R47H-*hTREM2* cDNA using Primers 5 and 6. Lane M = 1kb plus DNA ladder, Lanes 1-2 = R47H-*hTREM2*, Lanes 3-4 = CV-*hTREM2*, Lanes 5-6 = C57BL/6 control, Lanes 7-8 = H_2_O control. The PCR product size is 347 bp. **(g)** PCR detection of non-specific integration on 3’ end of the CV-*hTREM2* and R47H-*hTREM2* cDNA using Primers 7 and 8. Lane M = 1kb plus DNA ladder, Lanes 1-2 = R47H-*hTREM2*, Lanes 3-4 = CV-*hTREM2*, Lanes 5-6 = C57BL/6 control, Lanes 7-8 = H_2_O control. The PCR product size is 398 bp.

**Extended Data Figure 5.**
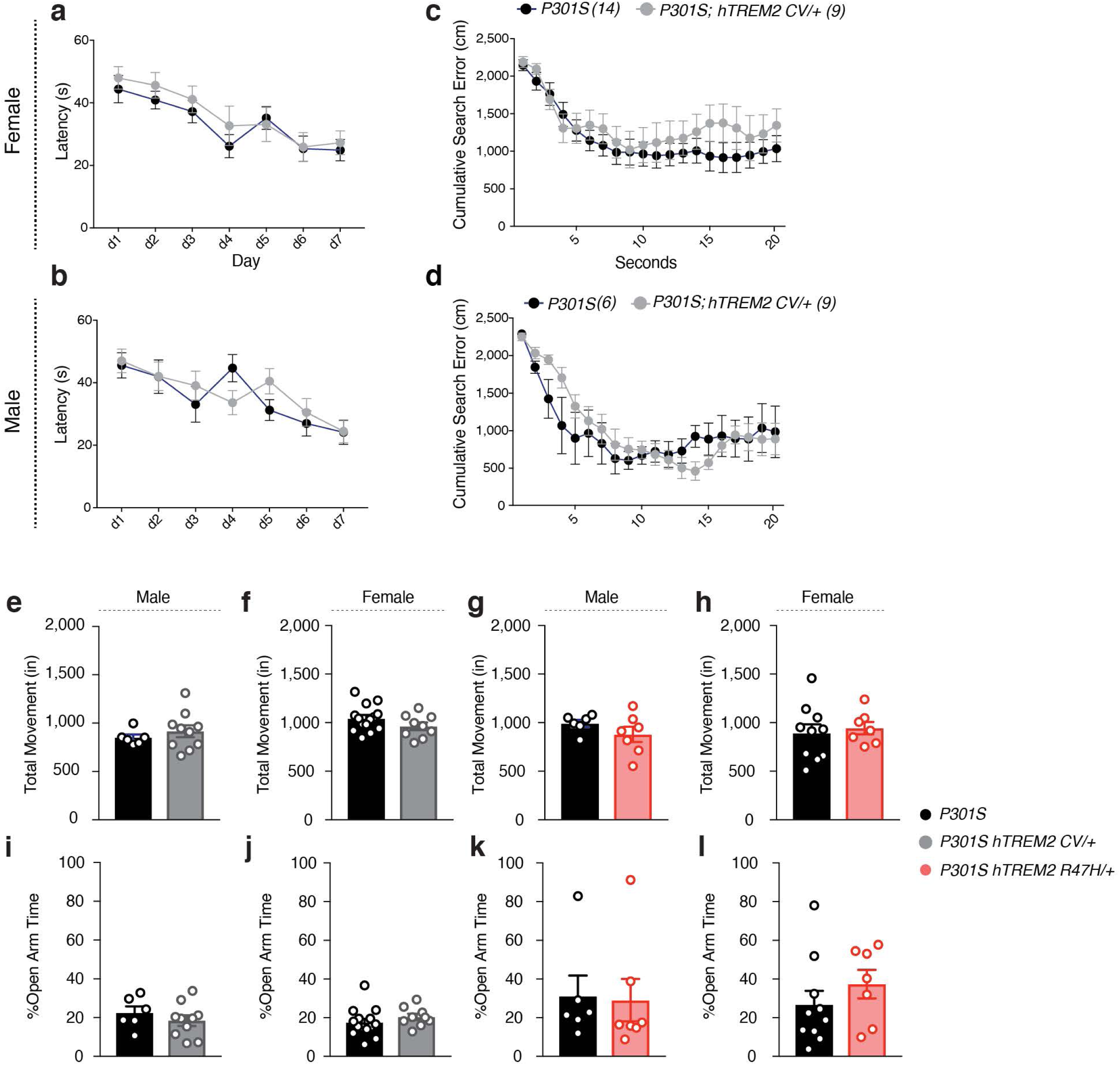
WT-*hTREM2* Tauopathy Mice Behave Similar to *mTrem2^+/+^* Tauopathy Mice (Related to Fig. 3) **(a-d)** Latency to reach the platform during hidden trials (d1-d7) and cumulative search errors during the 72-hr probe trial for female (**a,c**) and male (**b,d)** P301S and P301S *hTREM2^CV/+^* mice. STATA mixed-effects modeling for (**a,b)**. Two-tailed Mann-Whitney U-test of area under the curve for (**c,d)**. **(e-h)** Open-field assessment of overall activity of 7–9-month-old P301S *hTREM2^CV/+^* male (**e**) and female (**f**) and P301S *hTREM2^R47H/+^* male (**g**) and female (**h**) mice and littermate P301S controls. Two-tailed Student’s t-test for n>8, two-tailed Mann-Whitney U-test for n<8. **(i-l)** Elevated plus maze assessment of anxiety levels in 7–9-month-old *hTREM2^CV/+^* male (**i**) and female (**j**) mice and *hTREM2^R47H/+^* male (**k**) and female (**l**) mice and littermate P301S controls. Two-tailed Student’s t-test for n>8, two-tailed Mann-Whitney U-test for n<8.

**Extended Data Figure 6.**
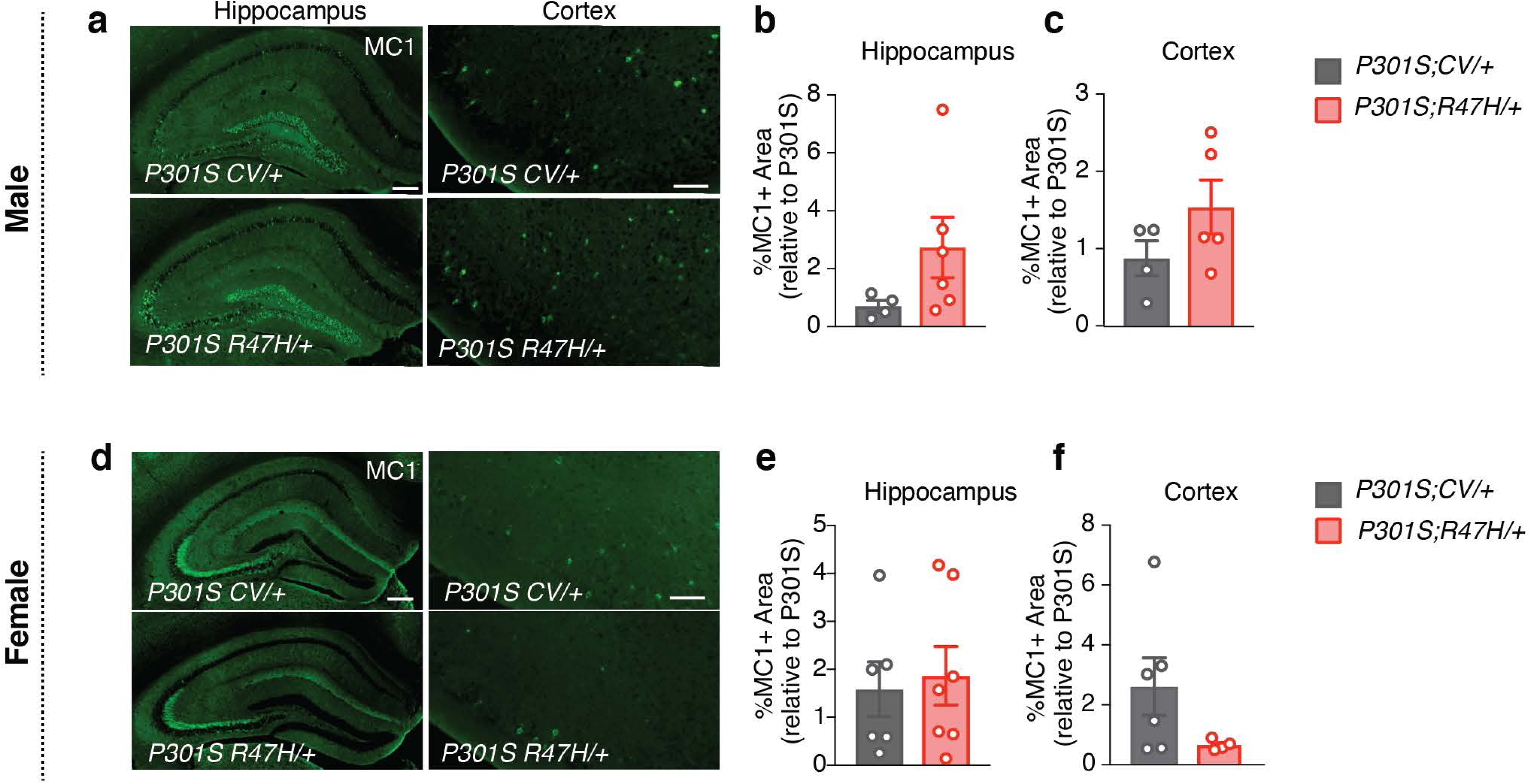
R47H-*hTREM2* Does not Alter Tau Pathology Load (Related to Fig. 3) Immunohistochemical analysis of 8–9-month-old mice. Each circle in bar graphs represents the average of 8–10 sections per mouse. Values are normalized to line-specific, sex-matched P301S *mTrem2*^+/+^ littermate controls. **(a,d)** Representative images of MC1 immunostaining in the hippocampus and entorhinal cortex of male (a) and female (**d**) mice. **(b-c, e-f)** Percentage of MC1+ area in male hippocampus (**b**) and cortex (**c**) and female hippocampus (**e**) and cortex (**f**) of P301S *hTREM2^CV/+^* and P301S *hTREM2^R47H/+^* mice. Entire hippocampus and entire cortex were quantified. Two-tailed Mann-Whitney U-test. Values are mean ± SEM.

**Extended Data Figure 7.**
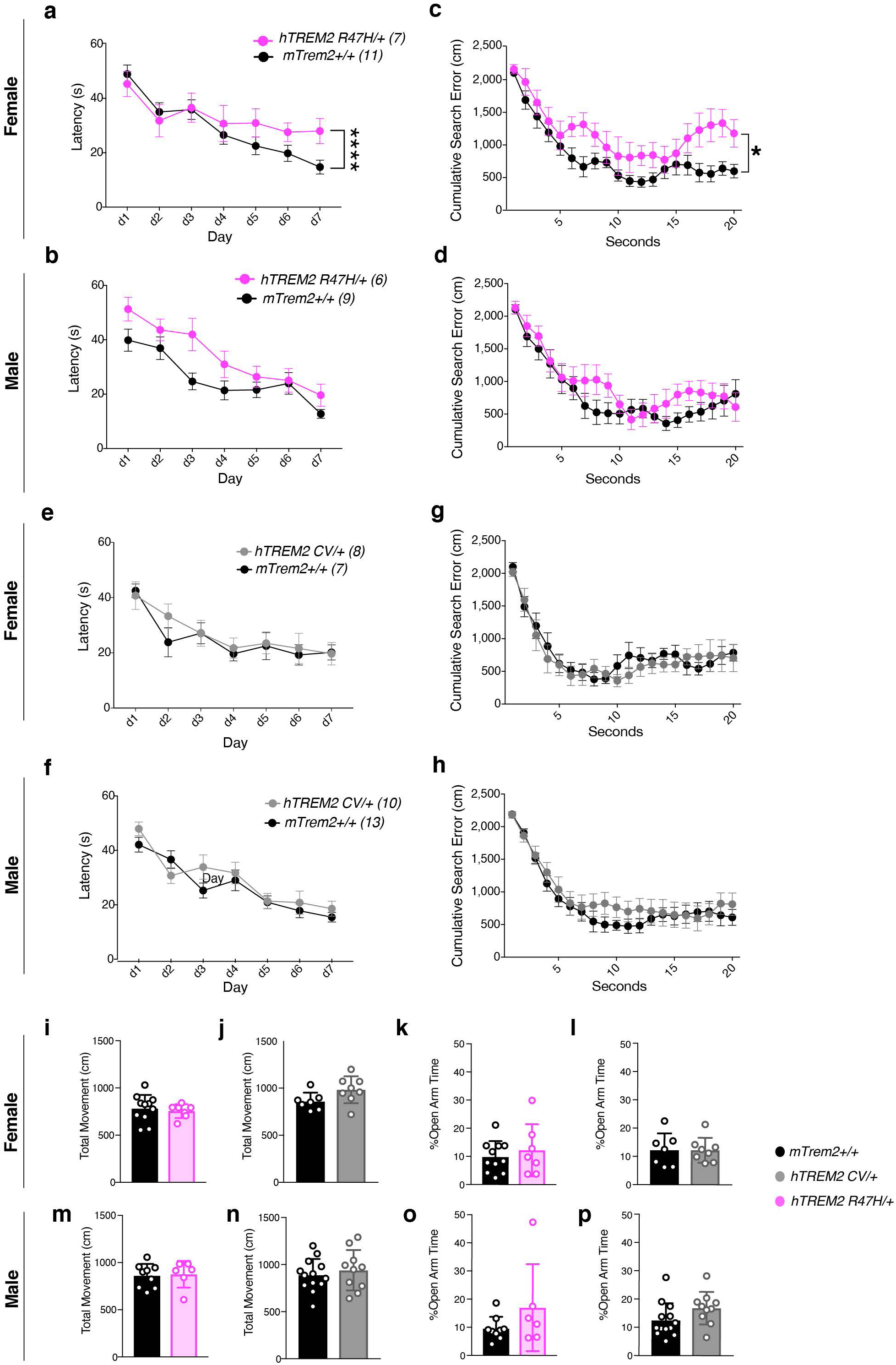
R47H-*hTREM2* Induces Spatial Learning and Memory Deficits in Female Mice (Related to Fig. 3) Morris water maze assessment of spatial learning and memory in 7–9-month-old *hTREM2^R47H/+^* and *hTREM2^CV/+^* mice and their littermate controls. **(a,b)** Latency to reach the platform during hidden trials (d1–d7) for *hTREM2^R47H/+^* females (**a**) and males (b) and their littermate sex-matched *mTrem2^+/+^* controls. ****p=0.0001, STATA mixed-effects modeling. **(c,d)** Cumulative search errors during the 24-hr probe trial for *hTREM2^R47H/+^* females (**c**) and males (**d**) and their littermate sex-matched *mTrem2^+/+^* controls. * U=13, p=0.0204, Two-tailed Mann-Whitney U-test of area under the curve. **(e,f)** Latency to reach the platform during hidden platform trials (d1–d7) for *hTREM2^CV/+^* females (**e**) and males (**f**) and their littermate sex-matched *mTrem2^+/+^* controls. STATA mixed-effects modeling. **(g,h)** Cumulative search errors during the 24-hr probe trial for *hTREM2^CV/+^* females (**g**) and males (**h**) and their littermate sex-matched *mTrem2^+/+^* controls. Two-tailed Mann-Whitney U-test of area under the curve. **(i-p)** Open-field assessment of overall activity **(i,j,m,n)** and elevated plus maze assessment of anxiety levels **(k,l,o,p)** of 7–9-month-old *mTrem2^+/+^, hTREM2^CV/+^*, and *hTREM2^R47H/+^* mice. Values are mean ± SEM.

**Extended Data Figure 8.**
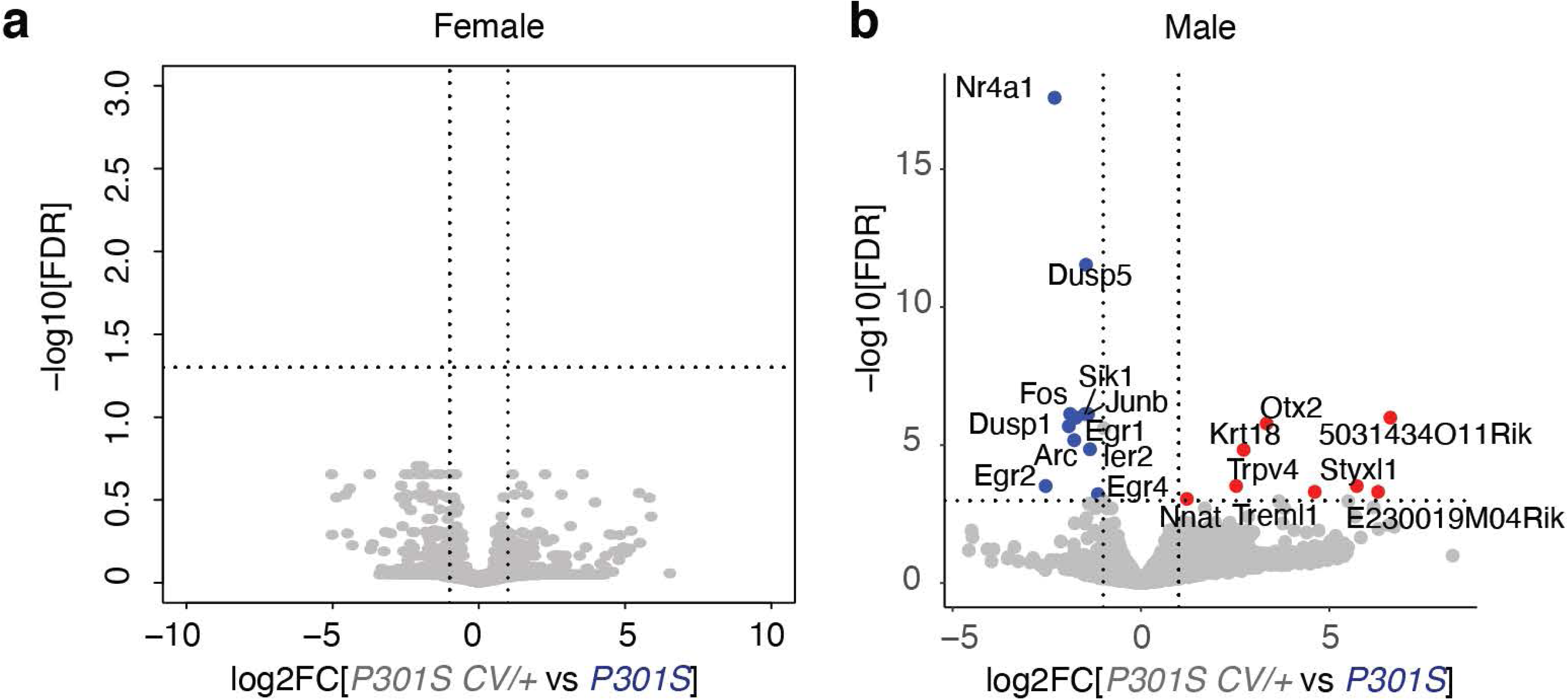
CV-hTREM2 has Similar Transcriptomic Profiles to mTrem2 (Related to Fig. 4). **(a-b)** Volcano plot of bulk RNA-seq data of hippocampal tissue from female (a) and male (b) P301S *hTREM2^CV/+^* and line-specific female and male P301S controls, respectively. Blue dots are genes with significantly higher normalized counts in P301S controls than in P301S *hTREM2^CV/+^* samples (11 mRNAs). Red dots are genes with significantly higher normalized counts in P301S *hTREM2^CV/+^* samples than P301S controls (7 mRNAs). Vertical dashed lines indicate log2FC ± 1. Horizontal dashed line indicates -log10(0.05). Wald test used. (n = 3 mice for P301S; n = 6 mice for P301S *hTREM2^CV/+^* for each sex).

**Extended Data Figure 9.**
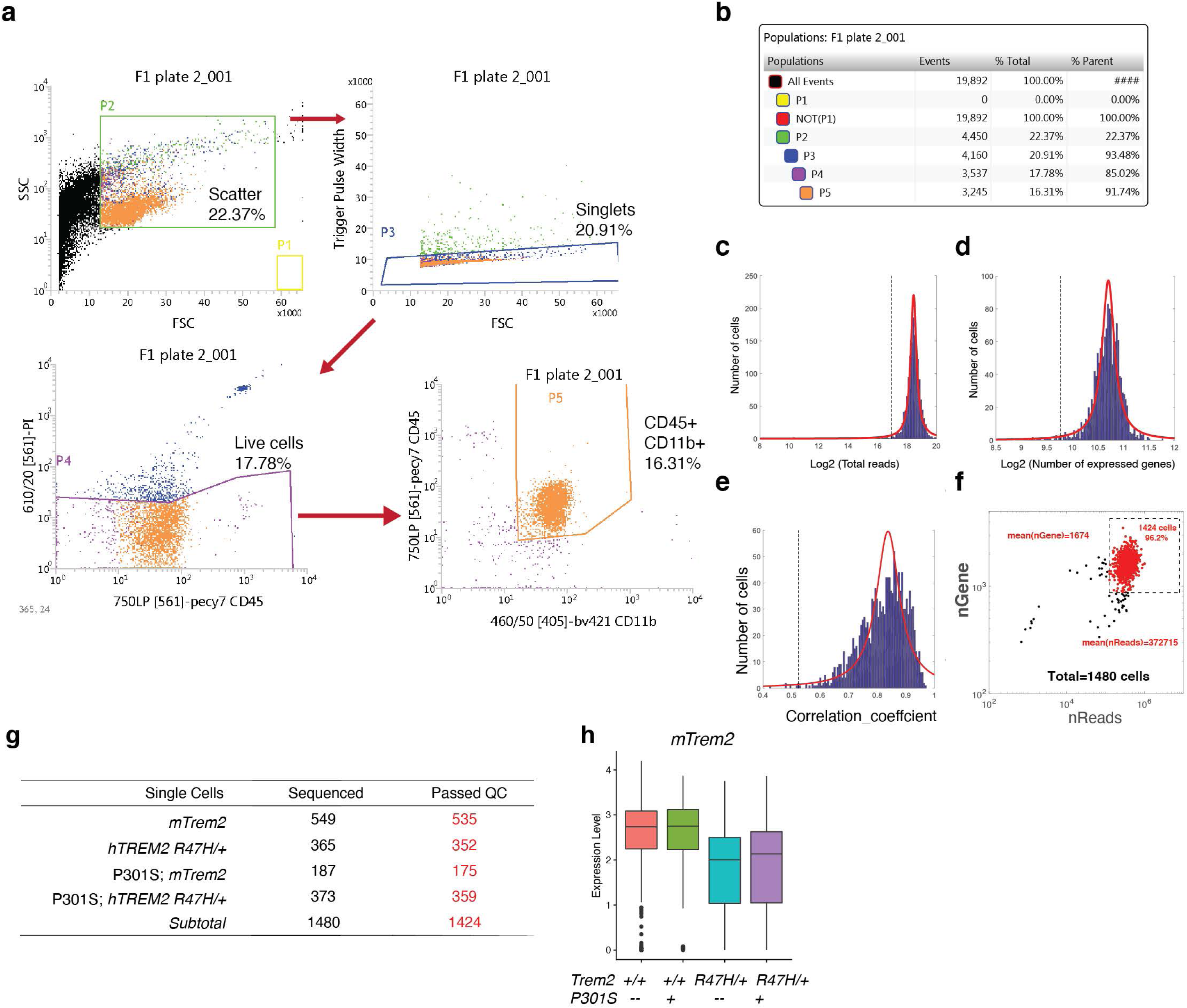
Single-Cell RNA-Seq of Brain CD45^+^;CD11b^+^ Cells (Related to Fig. 5) **(a)** Representative FACS plots showing gating strategies and cells sequenced. **(b)** Number of cells and proportions based on FACS from (**a**). **(c-e)** Quality control criteria for the single-cell sequencing data. Fitted curves for histograms of mapped reads (**c**), number of detected genes (**d**) and ERCC correlation coefficient **(e)** are labeled in red. Dashed lines are statistical cutoffs. Cells that passed all three criteria were retained for analysis. **(f)** Scatter plot highlighting cells that passed QC (red) among all cells sequenced. Each dot is a cell. **(g)** Summary table for the numbers of cells sequenced and cells that passed QC (red). 1424 cells were retained for analysis. **(h)** Normalized *mTrem2* expression level. Boxplot elements: center line, median; box limits, upper and lower quartiles; whiskers, 1.5x interquartile range; dots, outliers.

**Extended Data Figure 10.**
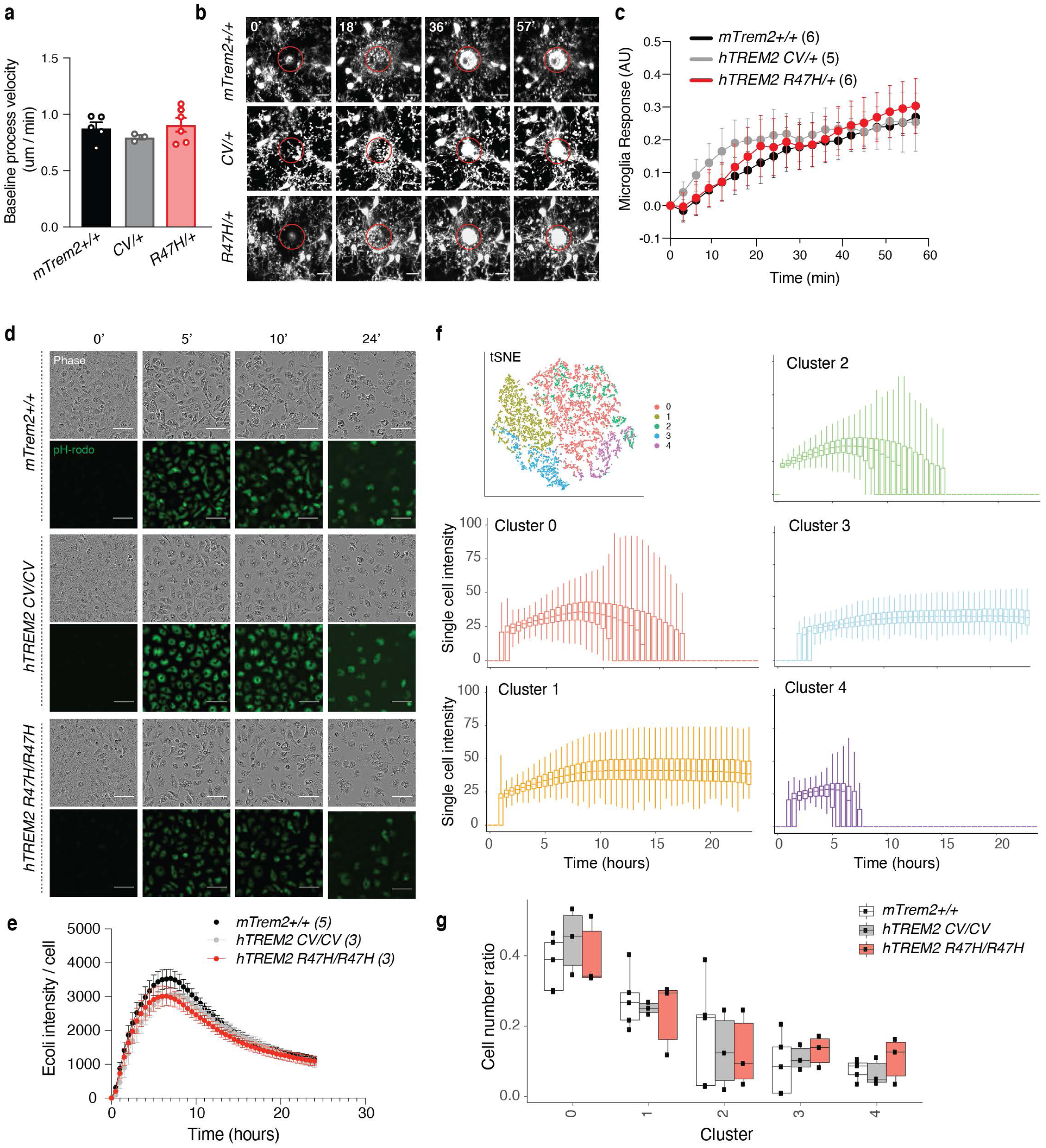
R47H-*hTREM2* Does Not Affect Microglial Injury Response or Phagocytosis (Related to Fig. 5) **(a)** Quantification of microglial process velocity in 12-17-month-old mice during 10 min of baseline recordings (n = 5 *mTrem2^+/+^,* n = 3 *hTREM2^CV/+^,* n = 6 *hTREM2^R47H/+^* mice, ∼25 processes/mouse). One-way ANOVA. **(b)** Representative images at 0, 18, 36, and 57 minutes after laser-induced injury. Red circle indicates region of injury. **(c)** Quantification of the normalized microglial response to focal laser-induced tissue injury for 60 min after injury. (n = 6 *mTrem2^+/+^,* n = 5 *hTREM2^CV/+^,* n = 6 *hTREM2^R47H/+^* mice). STATA mixed-effects modeling. Scale bars: hippocampus, 300 µm; cortex, 200 µm; hippocampal microglia, 20 µm, ablation, 15 µm. Values are mean ± SEM. All immunohistochemistry imaging and data are from one independent experiment. For *in vivo* imaging, one mouse was imaged per imaging session. **(d)** Representative images of primary microglia at 0, 5, 10, and 24 hours after incubation with pH-rodo conjugated *E. coli* substrates. Scale bar: 100 µm. **(e)** Bulk fluorescence intensity quantification of field of view normalized by cell number. (n = 5 mTrem2+/+, n = 3 *hTREM2^CV/+^,* n = 3 *hTREM2^R47H/+^* biologically independent experiments). **(f)** tSNE of single-cell quantification of fluorescence intensity of all cells analyzed and fluorescence intensity change over time plotted as boxplots for each time point for each cellular cluster. Boxplot elements: center line, median; box limits, upper and lower quartiles; whiskers, 1.5x interquartile range. **(g)** Boxplot of proportions of each genotype for each cell cluster determined in (f). Boxplot elements: center line, median; box limits, upper and lower quartiles; whiskers, 1.5x interquartile range; dots, individual experimental values. See also Supplementary Movie 1.

## SUPPLEMENTAL MOVIES AND TABLES

**Supplementary Movie 1. R47H-*hTREM2* Does Not Affect Microglial Response to Injury (Related to Extended Data Fig. 10)**

Video file of microglial responses of 12-17-month-old *Cx3cr1^GFP/+^ mTrem2^+/+^*, *hTREM2^CV/+^*, and *hTREM2^R47H/+^* mice to an acute laser-induced injury *in vivo* over a 60-min time period, played one after the other. Scale bar: 15 µm.

**Supplementary Table 1: Patient Clinical Information (Related to Figure 1)**

Table of characteristics pertaining to the 16 donors whose brain tissues were used for single-nuclei RNA-sequencing.

**Supplementary Table 2: Sequencing Information and Quality Control Metric for All AD Tissue Samples Sequenced (Related to Figure 1)**

Table of sequencing metrics for all 16 samples used for single-nuclei RNA-sequencing.

**Supplementary Table 3: DEGs Induced by the R47H Mutation in Each Cell Type and Sex in AD Patient Brains (Related to Figure 1)**

Table summarizing differential expression analysis between R47H-AD cells vs CV-AD cells for each brain cell type and each patient sex.

**Supplementary Table 4: Markers for the Microglia Subpopulations Isolated from AD Tissues (Related to Figure 2)**

Table summarizing differential expression analysis between a specific microglial cluster versus the homeostatic microglia cluster 1.

**Supplementary Table 5. DEGs Induced by R47H-hTREM2 in Female Tauopathy Mice (Related to Figure 4)**

Table of differential expression analysis between female P301S hTREM2R47H/+ mice and line-specific female P301S controls.

**Supplementary Table 6: WGCNA Brown and Cyan Module Genes (Related to Figure 4)**

Table of genes in the brown and cyan module of the weighted gene co-expression network analysis of bulk RNA-sequencing data from female P301S *hTREM2^WT/+^,* P301S *hTREM2^R47H/+^* mice and line-specific female P301S controls.

**Supplementary Table 7: Marker Genes for Cluster 2 of Microglia from Single-Cell RNA-Seq of Female Mice (Related to Figure 5)**

Table of differential expression analysis between microglia cluster 2 vs cluster 1 from single-cell RNA-seq of 8-month-old female *hTREM2^R47H/+^*, P301S *hTREM2^R47H/+^*, and their line-specific female controls.

**Supplementary Table 8: DEGs Induced in R47H/R47H Microglia in Response to Tau Fibrils (Related to Figure 6).**

Table of differential expression analysis between *hTREM2^R47H/R47H^* vs. *mTrem2^+/+^* primary microglia treated with tau fibrils.

**Supplementary Table 9: DEGs between R47H/R47H Microglia Treated with MK-2206 versus Vehicle in Response to Tau Fibrils (Related to Figure 6).**

Table of differential expression analysis between *hTREM2^R47H/R47H^* primary microglia treated with Akt-inhibitor MK-2206 vs. *hTREM2^R47H/R47H^* primary microglia treated with vehicle, both in the presence of tau fibrils.

