## Supplementary figures and images for "AD-linked R47H-*TREM2* mutation induces disease-enhancing proinflammatory microglial states in mice and humans"

### Source Data

Source Data for Figure 3.

#### Full Blots

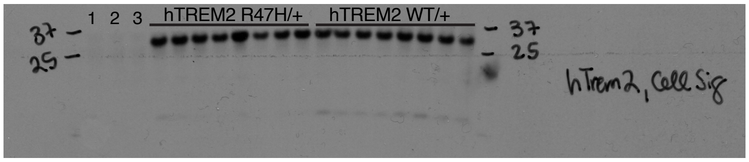


1 = mTrem2 KO
2-3 = mTrem2+/+

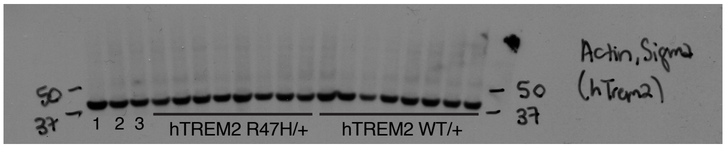
